# Indigenous Yanomami gut microbiota protects against diet-induced metabolic dysfunction

**DOI:** 10.64898/2026.10.28.762182

**Authors:** Marie Thorp, Lucas Amoroso Lopes de Carvalho, Maarten Ciers, Gillian Blancke, Vanessa Andries, Maude Jans, Sergio García Sánchez, Britta De Pessemier, Chris Callewaert, David Good, Geert van Loo, Emma Allen-Vercoe, Luciana Campos Paulino, Lars Vereecke

**Affiliations:** VIB Center for Inflammation Research, Ghent, Belgium; Department of Internal Medicine and Pediatrics, Ghent University, Ghent, Belgium; Center for Natural Sciences and Humanities, Federal University of ABC, Santo André, São Paulo, Brazil; Department of Biomedical Molecular Biology, Ghent University, Ghent, Belgium; Center for Microbial Ecology and Technology, Ghent University, Ghent, Belgium; Department of Molecular and Cellular Biology, University of Guelph, Guelph, Ontario, Canada

**Keywords:** gut microbiota, industrialization/urbanization, metabolic disease, germ-free mice, microbiota transplantation

## Abstract

Industrialization profoundly alters the human gut microbiota, reducing microbial diversity and eliminating taxa prevalent in non-industrialized populations. These microbiota alterations accompanying industrialization might contribute to the increasing prevalence of obesity and other chronic inflammatory diseases, yet direct experimental evidence remains lacking. We compared the gut microbiota of Indigenous Brazilian Yanomami and urban Belgian donors and transplanted these into germ-free (GF) mice subjected to high-fat diet (HFD). The Yanomami microbiota exhibited greater richness and distinct composition enriched for VANISH (Volatile and/or Associated Negatively with Industrialized Societies of Humans) taxa. Transplantation of Yanomami microbiota significantly attenuated HFD-induced metabolic dysfunction, reducing weight gain, diabetes incidence, and promoting adipose and hepatic Th2 immunity. Yanomami-derived microbial communities displayed enhanced ecological stability and enrichment for short-chain fatty acid-associated taxa and metabolic functions post-HFD. Our findings demonstrate that the microbiota from low-urbanization populations confer protection against diet-induced metabolic disease, highlighting ecological resilience and specific microbial functions as potential targets for microbiota-inspired therapeutics.

## Introduction

The Yanomami are an Indigenous people whose communities extend across northern Brazil and southern Venezuela. The communities participating in this study, located along the Marauiá River in Amazonas, Brazil, continue to rely on small-scale swidden horticulture, fishing, hunting, gathering, and locally produced foods. Their diet is rich in fibre, consisting primarily of fruits and vegetables (including plantains and cassava), wild caught meat and fish, and is inherently low in fat and salt^1,2^. Non-communicable diseases common in urban industrialized societies – such as obesity, hypertension, and chronic inflammatory or autoimmune diseases – are reported to be virtually absent among the Yanomami communities studied so far^3–7^.

Historically, microbiome research has focused mainly on urban industrialized populations (often referred to as “Western”), resulting in an underrepresentation of Indigenous and non-urbanized populations^8^. However, Western lifestyle factors – including diet, sanitation, modern medicine, and widespread antibiotic use – have well documented impacts on the gut microbiota and are associated with reduced microbial diversity^9–11^. Therefore, over the past decade, researchers have increasingly turned to characterizing the microbiomes of non-urbanized populations, such as the Yanomami, to broaden our understanding of human microbiome diversity across populations with markedly different degrees of urbanization and industrialization.

Multiple studies have reported unprecedented faecal bacterial diversity in the gut microbiomes of Yanomami individuals^2,12^. Also the Hadza population in Tanzania harbour a highly diverse and seasonally varying gut microbiome, with many recovered genomes absent from current reference databases^13,14^. This study introduced the concept of “VANISH” microbes (Volatile and/or Associated Negatively with Industrialized Societies of Humans) consistently found in the Hadza gut microbiomes but absent in industrialized populations. In contrast, "BloSSUM" microbes (Bloom or Selected in Societies of Urbanization/Modernization) are abundant in industrialized populations but absent in the Hadza community. Indeed, enrichment of VANISH taxa in traditional populations has been reported across many populations spanning sub-Saharan Africa^15–19^, South America^12,20,21^, Oceania^22^, and Southeast Asia^23–25^, underscoring the profound impact of industrialization on the human gut microbiome.

The gut microbiome is known to play a fundamental role in immune maturation and host physiology, and has been implicated in the development of metabolic disease. Indeed, comparative studies have reported compositional differences between lean and obese phenotypes, most classically an increase in Firmicutes/Bacteroidetes ratio (renamed Bacillota/Bacteroidota) in obese humans and genetically obese mice^26,27^. Subsequent metagenomic analysis showed that obesity is associated with reduced overall microbial gene richness and altered representation of metabolic pathways, including those involved in energy harvest from dietary substrates^28,29^. Further causal evidence for the effect of specific taxa on obesity has come from gnotobiotic mouse models. Colonization of germ-free (GF) mice with a conventional specific pathogen free (SPF) microbiota increases adiposity despite reduced food intake, establishing the gut microbiota as a regulator of host energy storage^30^. This was extended to human-derived microbiota: transplantation of faecal communities from obesity-discordant twin pairs into GF mice transmitted the donor’s adiposity phenotype in a diet-dependent manner^31^. Furthermore, several studies have demonstrated protective effects of specific bacterial taxa, including *Phascolarctobacterium faecium*^32,33^, *Prevotella copri*^34^, *Limosilactobacillus reuteri*^35^, and *Akkermansia muciniphila*^36^ against high-fat diet (HFD)-induced obesity and associated metabolic dysfunction. The latter has since been translated to humans: a randomized, placebo-controlled pilot trial showed that oral *A. muciniphila* supplementation improved insulin sensitivity and reduced circulating cholesterol in overweight or obese volunteers^37^, providing clinical proof-of-concept that a single bacterial taxon can causally modulate host metabolic parameters.

The role of an Indigenous human microbiota in the development of metabolic disorders remains unexplored. Therefore, this study investigates compositional differences between urban industrialized (Belgian) and Indigenous Yanomami gut microbiota and evaluates the functional consequences of these distinct microbial communities in the context of HFD-induced obesity. Our results demonstrate that the Yanomami gut microbiota is more diverse and compositionally distinct from the Belgian microbiota, and that its transfer into GF mice confers protection against multiple features of HFD-induced metabolic disease, associated with a more diet-resilient, short-chain-fatty-acid–oriented microbial community.

## Results

### Yanomami gut microbiota exhibit a distinct low-urbanization signature

Faecal samples were collected from the Indigenous Yanomami participants living in communities along the Rio Marauiá in northern Brazil. As an urban industrialized comparison cohort, faecal samples were collected from Belgian individuals (Fig. 1a). Participants had no diagnosed metabolic, inflammatory or immune-mediated diseases, and showed no clinical signs or symptoms of acute or chronic illness or infection at the time of sampling. They had not received antibiotics within three months preceding sample collection. In both Belgian (n = 39) and Yanomami (n = 78) cohorts, samples were collected from male and female donors, between the ages of 2 and 64 years old (Fig. 1b). During sampling, information was collected on the lifestyle and diet of participants, consistent with a high fibre, low fat dietary pattern of Yanomami participants, with high consumption of cassava flour, plantain, legumes, fish and occasional meat (Suppl. Fig. 1a). A companion metagenomic study provides a detailed characterization of microbiome composition and functional potential across these Yanomami communities in relation to varying degrees of urban influence (de Carvalho et al., *in prep*).

**Fig. 1.**
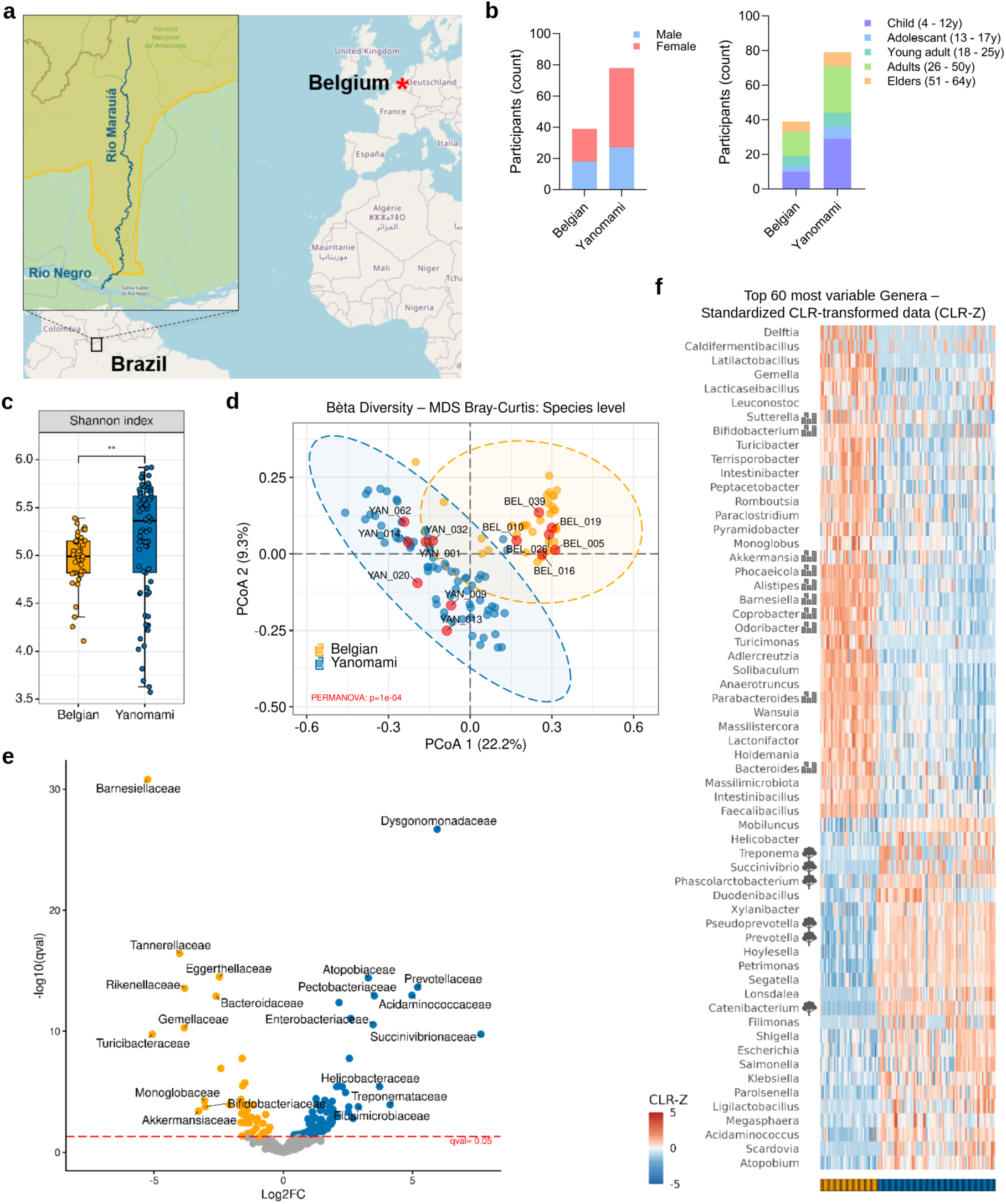
Sample collection and metagenomic profiling of Belgian and Yanomami gut microbiota. **a,** Sampling locations along the Rio Marauia in the Brazilian Yanomami territory (yellow) and Belgium (source: OpenStreetMap) **b,** Sex (left) and age (right) distribution of Belgian (n=39) and Yanomami (n=78) human participants. **c**, Distribution of gut microbiota alpha diversity in Belgian and Yanomami cohort. Shannon index values were calculated at the taxonomic feature level from contig-based classifications, using the best taxonomic assignment up to the species level. Statistical significance between groups was assessed using the Wilcoxon test (P = 0.001; W = 962; Hodges-Lehmann estimate = -0.34). **d**, Principal coordinate analysis (PCoA) of Belgian vs. Yanomami gut microbiota profiles at species level showing strong clustering by donor cohort (Bray-Curtis dissimilarity metric; PERMANOVA: P = 0.0001). Red dots indicate donors used for in vivo experiments. **e**, Volcano plot of differentially abundant taxa (Yanomami vs. Belgian stool) at the family level. Differential abundance analysis was performed using MaAsLin2. Statistical significance was defined at qval < 0.05. The 20 taxa with the largest absolute effect sizes (10 positively and 10 negatively associated) are labelled. **f**, Heatmap of most variable genera when comparing Belgian and Yanomami metagenomic profiles. For each genus, counts were CLR-transformed to account for compositionality and then Z-scaled across samples (CLR-Z). VANISH (Volatile and/or Associated Negatively with Industrialized Societies of Humans) taxa are indicated by a tree icon, while BloSSUM (Bloom or Selected in Societies of Urbanization/Modernization) taxa are indicated by an urban icon.

Taxonomic alpha diversity, assessed using the Shannon index, was significantly higher in the Yanomami gut microbiota (Fig. 1c). This increased alpha diversity is accompanied by greater species richness, whereas the Gini-Simpson index did not differ between groups (Suppl. Fig. 1b). This pattern is consistent with an expansion of low-abundance taxa contributing to the higher diversity observed in the Yanomami microbiota. Furthermore, community compositional profiles of Belgian and Yanomami gut microbiota are distinct, as shown by principal coordinate analysis (PCoA) at species level (Bray-Curtis dissimilarity metric; Fig. 1d). To enable functional comparison of representative microbiota, we used Bray-Curtis dissimilarity to exclude donor samples at extreme ends of each cohort, while accounting for sample availability and covariate matching, for subsequent human microbiota-associated (HMA) mouse experiments (indicated in red on Fig. 1d, see further).

Differential abundance analysis revealed extensive taxonomic differences between the two cohorts, with 105 microbial families enriched in the Yanomami microbiota and 45 in the Belgian microbiota (Fig. 1e). Consistent with previous reports, Yanomami microbiota were enriched for VANISH-associated families^13^, including Prevotellaceae, Succinivibrionaceae and Treponemataceae, while BloSSUM-associated families, such as Bacteroidaceae, Rikenellaceae and Akkermansiaceae are more abundant in the Belgian gut microbiota (Fig. 1e, Suppl. Fig. 1c). This observation translates to the genus level, where we see an increased relative abundance of *Segatella* (part of Prevotellaceae) and a decrease of *Bacteroides* (part of Bacteroidaceae, Suppl. Fig. 1d). Indeed, this separation of VANISH and BloSSUM taxa is clear when looking at the top 60 most variable genera (Fig. 1f), with VANISH genera increased in the Yanomami gut microbiota alone (indicated with tree icon), and BloSSUM genera increased in the Belgian group (indicated with urban icon). Furthermore, several genera enriched in the Yanomami microbiota — including *Helicobacter*, *Mobiluncus*, *Megasphaera*, *Atopobium*, *Scardovia*, *Ligilactobacillus*, *Petrimonas*, *Duodenibacillus*, and *Parolsenella* — have not yet been classified as VANISH taxa but showed no evidence of industrialized-population enrichment elsewhere in literature, positioning them as candidate novel VANISH-associated taxa warranting independent validation in additional non-industrialized cohorts.

The greater diversity and enrichment of VANISH taxa observed here using read-level taxonomic profiling were also recovered in a companion study using a metagenome assembly–based strategy (de Carvalho et al., *in prep*). Assembly-based functional profiling further showed that Yanomami microbiota carried a distinct encoded metabolic repertoire, including consistent enrichment of Kyoto Encyclopedia of Genes and Genomes (KEGG) butanoate metabolism across all Yanomami populations studied. Their genus-level association networks were substantially more connected and more robust to simulated targeted node removal than the Belgian network, with both properties increasing as urban influence declined. These findings establish short-chain fatty acid (SCFA)-associated metabolic potential and a highly connected association-network architecture as prominent features of the Yanomami donors microbiota.

### Transplantation of a Yanomami gut microbiota reduces sensitivity to high-fat diet-induced metabolic dysfunction

The markedly lower prevalence of obesity and metabolic syndrome in low-urbanization populations has raised the possibility that differences in gut microbiota composition contribute to improved metabolic health. To validate this, we generated Belgian- or Yanomami-HMA mice (BEL_026 vs. YAN_013, Table 1), which were challenged with a HFD for 12 weeks (Fig. 2a).

**Fig. 2.**
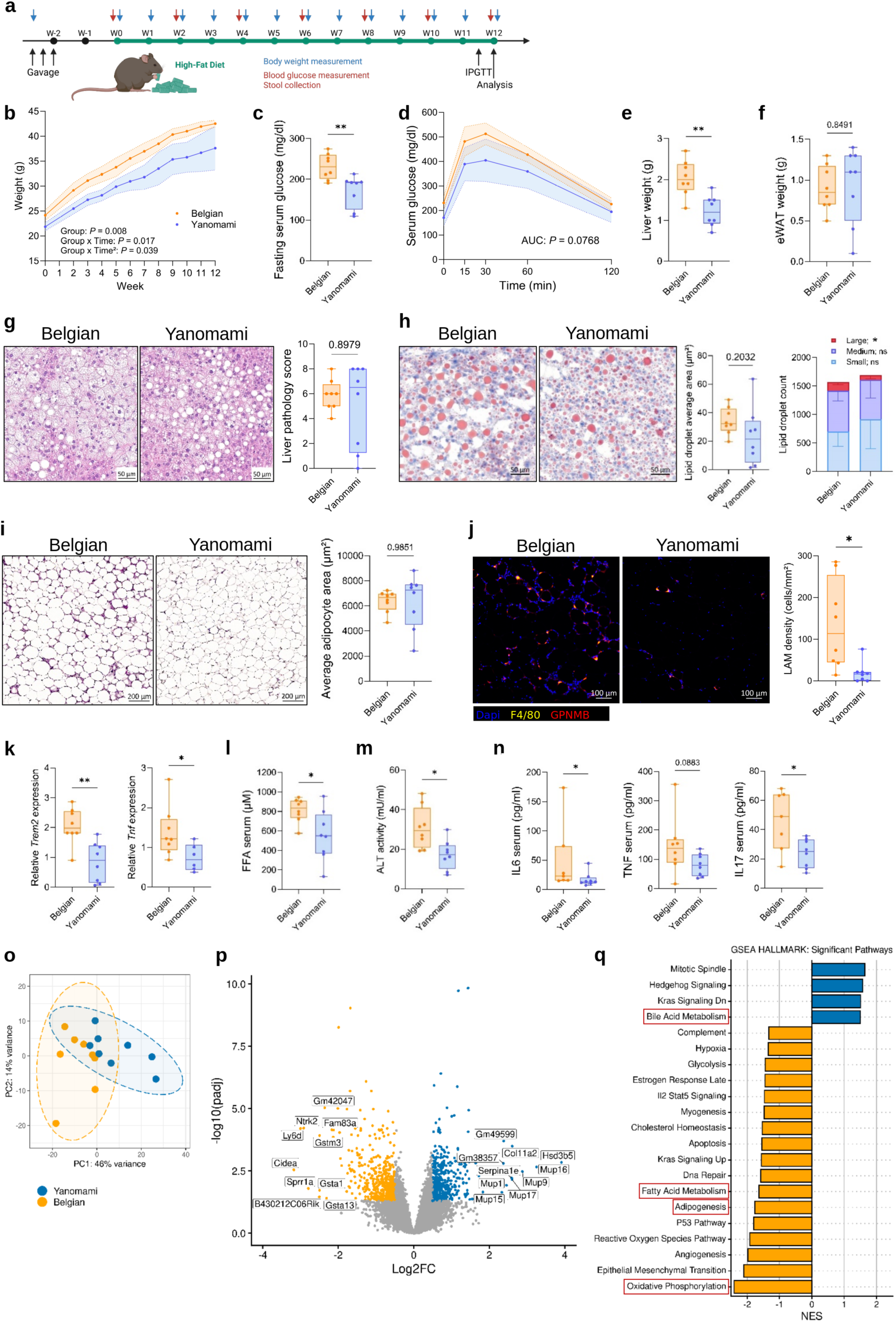
A Yanomami gut microbiota reduces sensitivity to high-fat diet-induced metabolic pathology. **a,** Experimental set-up of high-fat diet (HFD)-induced obesity mouse model. Image made with Biorender. **b**, Weight follow-up over the course of HFD administration (n = 8 per group). Data are represented as mean ± 95% CI. Data were analysed using a linear mixed-effects model fitted by restricted maximum likelihood (REML) with fixed effects for Time, Group, Group x Time and Group x Time² interactions, and random effects for mouse ID. Corresponding P values are indicated. **c**, Fasting serum glucose measurements of Belgian vs. Yanomami mice (16h fasting period) following 12 weeks of HFD. Significance was determined using a Welch’s t test (t(13,35) = 3.395; P = 0.0046; mean difference −60.25; 95% CI −98.49 to −22.01). **d**, Intraperitoneal glucose tolerance test (IPGTT). Mice were fasted for 16h and injected peritoneally with glucose (10%, 1g kg^-1^ body weight). Blood glucose levels were measured at the indicated time points. Data are shown as mean ± 95% CI. Significance of area under the curve (AUC) was determined using a Welch’s t test (t(10,45) = 1.96; P = 0.077; mean difference −8.263; 95% CI −17.589 to 1.063). **e-f**, Liver (e) and eWAT (f) weight measurement of Belgian vs. Yanomami mice following 12 weeks of HFD. Significance was determined using a Welch’s t test (liver: t(13,70) = 3.980; P = 0.0014; mean difference −0.8125; 95% CI −1.251 to −0.3738; eWAT: t(11,37) = 0.1946; P = 0.8491; mean difference 0.0375; 95% CI −0.3849 to 0.4599). **g**, Representative liver histology images (H&E, left) of Belgian vs. Yanomami mice following 12 weeks of HFD, with associated liver pathological scoring (right). Significance was determined using a Mann-Whitney test (P = 0.8979). **h**, Representative liver histology images (Oil Red O, left) of Belgian vs. Yanomami mice following 12 weeks of HFD, with associated measurement of average lipid droplet area (stained red on image, middle). Each datapoint represents the average of measured lipid droplets in 3 regions of a liver section of one mouse. Significance was determined using a Welch’s t test (t(9,913) = 1.362; P = 0.2032; mean difference −10.89; 95% CI −28.73 to 6.944). Lipid droplets were categorized as small/medium/large and counted (right). Significance of large lipid droplet count was determined using a Welch’s t test (t(13,06) = 2.322; P = 0.0370; mean difference −66.91; 95% CI −129 to -4.683). **i**, Representative eWAT histology images (H&E, left) of Belgian vs. Yanomami mice following 12 weeks of HFD, with associated measurement of adipocyte area (right). Each datapoint represents the average of measured adipocytes in an eWAT section of one mouse. Significance was determined using a Welch’s t test (t(9,193) = 0.01913; P = 0.9851; mean difference −15.66; 95% CI −1862 to 1830). **j,** Representative eWAT immunofluorescent images (blue: Dapi, yellow: F4/80, red: GPNMB) of Belgian vs. Yanomami mice following 12 weeks of HFD, with associated measurement of LAM density (F4/80+ GPNMB+ cells, right). Each datapoint represents the average of measured LAM density in 3 regions of an eWAT section of one mouse. Significance was determined using a Welch’s t test (t(7,739) = 2.979; P = 0.0183; mean difference −115.6; 95% CI -205.5 to -25.58). **k**, Gene expression analysis of Trem2 and Tnf in eWAT tissue of mice following 12 weeks of HFD. Gene expression was quantified relative to Ywhaz and Rplp0 expression levels. Significance was determined using a Mann-Whitney test (Trem2: Hodges-Lehmann difference = −1.144, Mann-Whitney U = 4, exact two-tailed P = 0.0017; Tnf: Hodges-Lehmann difference = −0.5496, Mann-Whitney U = 7, exact two-tailed P = 0.0293). **l**, Free fatty acid (FFA) levels measured in serum of mice following 12 weeks of HFD (non-fasting, 120min post glucose injection for IPGTT). Significance was determined using a Welch’s t test (t(9,966) = 2.572, P = 0.0279; mean difference −262.6, 95% CI −490.2 to −34.97). **m**, Serum ALT activity measured in serum of mice following 12 weeks of HFD (non-fasting, 120min post glucose injection for IPGTT). Significance was determined using a Mann-Whitney test (Hodges-Lehmann difference = −12.29; Mann-Whitney U = 8, n = 8 per group, exact two-tailed P = 0.0104). **n**, Inflammatory cytokines measured in serum of mice following 12 weeks of HFD (non-fasting, 120min post glucose injection for IPGTT). Significance was determined using a Mann-Whitney test (IL6: Mann-Whitney U = 8, exact two-tailed P = 0.0205; TNF: Mann-Whitney U = 15.5, exact two-tailed P = 0.0883; IL17: Mann-Whitney U = 9, exact two-tailed P = 0.0289). **o-q**, Bulk mRNA sequencing of liver tissue from Belgian vs. Yanomami mice following 12 weeks of HFD. **o**, Principal component analysis (PCA) of variance-stabilized (VST) read counts across all samples. Each point represents one biological replicate, with colours indicating experimental groups. Percentages on the axes indicate the proportion of variance explained by each principal component. **p,** Volcano plot showing the top 20 differentially expressed genes between Yanomami (blue)- and Belgian (yellow)-HMA liver samples. **q**, Gene set enrichment analysis (GSEA) of MSigDB Hallmark gene sets. Bar plot showing all significantly enriched Hallmark gene sets (blue: enriched in Yanomami-HMA liver; yellow: enriched in Belgian-HMA liver). NES: normalized enrichment score. LAM: Lipid-Associated Macrophage; *: P < 0.05; **: P < 0.01

**Table 1.** Overview of human donor samples used during high-fat diet (HFD)-induced metabolic disease experiments. Sex, age and BMI status of human donors is listed between brackets. nw: normal weight, BMI = 18.5 – 25; ow: overweight, BMI = 25-30; ob: obese, BMI > 30; B: Belgian; Y: Yanomami.

| Belgian donor | Yanomami donor | Mice/donor |
| --- | --- | --- |
| BEL_051 (male, 7, nw) | YAN_009 (male, 7, nw) | 8 (B) + 8 (Y) |
| BEL_026 (female, 24, nw) | YAN_013 (female, 22, nw) | 8 (B) + 8 (Y) |
| BEL_016 (male, 30, nw) | YAN_032 (male, 36, ow) | 3 (B) + 3 (Y) |
| BEL_039 (female, 23, nw) | YAN_014 (female, 16, nw) | 3 (B) + 3 (Y) |
| BEL_005 (female, 8, nw) | YAN_062 (female, 4, nw) | 3 (B) + 3 (Y) |
| BEL_019 (female, 25, ob) | YAN_001 (female, 37, ow) | 3 (B) + 3 (Y) |
| BEL_010 (male, 4, nw) | YAN_020 (female, 4, nw) | 4 (B) + 3 (Y) |

Transplantation of a Yanomami microbiota markedly reduced susceptibility to HFD-induced metabolic dysfunction. At the start of the feeding period, Yanomami-HMA mice had a significantly lower body weight than control mice (estimated difference = −2.43 g, *P* = 0.008). Body weight changed nonlinearly over time, and significant Group × Time (*P* = 0.017) and Group × Time² (*P* = 0.039) interactions indicated that the groups followed different body-weight trajectories over time (Fig. 2b). In addition to differences in body weight trajectories, Yanomami-HMA mice exhibited improved glucose homeostasis, reflected by significantly lower fasting glucose concentrations (Fig. 2c) and a trend towards improved glucose tolerance during intraperitoneal glucose tolerance testing (IPGTT, Fig. 2d). Although epididymal white adipose tissue (eWAT) weight was comparable between groups (Fig. 2f), Yanomami-HMA mice displayed significantly lower liver weights (Fig. 2e), suggesting reduced hepatic metabolic pathology.

To determine whether the improved metabolic phenotype of Yanomami-HMA mice was accompanied by reduced hepatic pathology, liver sections were evaluated by haematoxylin and eosin (H&E) staining. Both Belgian- and Yanomami-HMA mice developed hepatocellular hypertrophy and micro- and macrovesicular steatosis following HFD, with no significant differences in histopathological score (Fig. 2g). We then performed a more detailed quantification of hepatic lipid accumulation using Oil Red O (ORO) staining. While the mean lipid droplet area was reduced in Yanomami-HMA mice without reaching statistical significance (Fig. 2h), stratification of lipid droplets by size revealed a significant reduction in large lipid droplets (defined as the 5% largest lipid droplets of all measurements) compared to Belgian-HMA mice. Consistently, analysis of lipid droplet size distribution demonstrated a shift towards smaller lipid droplets in Yanomami-HMA livers, whereas Belgian-HMA mice accumulated large lipid droplets (Suppl. Fig. 2a). Together, these findings indicate that transplantation of a Yanomami microbiota alters hepatic lipid storage, limiting accumulation of large lipid droplets despite comparable histopathology.

We next examined whether transplantation of a Yanomami microbiota also altered adipose tissue morphology and inflammation. H&E staining of eWAT revealed no significant differences in adipocyte size between Belgian- and Yanomami-HMA mice (Fig. 2i), indicating that the reduced body weight of Yanomami-HMA mice was not accompanied by overt changes in adipocyte hypertrophy. In contrast, Belgian-HMA mice displayed a visibly greater abundance of crown-like structures (CLS), which are formed by lipid-associated macrophages (LAMs) surrounding dead or dying adipocytes. To quantify this observation, eWAT sections were immunostained for LAM markers F4/80 and GPNMB (Fig. 2j), confirming reduced accumulation of LAMs in Yanomami-HMA mice.

To determine whether this reduced accumulation of LAMs was accompanied by altered inflammatory signalling, we profiled gene expression in eWAT by RT–qPCR. Consistent with the histological findings, Yanomami-HMA mice exhibited significantly lower expression of *Trem2* and *Tnf* confirming reduced LAM infiltration and associated inflammation (Fig. 2k). Expression of other inflammatory markers, including *F4/80, Mcp1, Il-6, Il-1β, Il-10* was not significantly altered (Suppl. Fig. 2b, left). Furthermore, we evaluated gene expression of several metabolic markers and adipokines in eWAT (Suppl. Fig. 2b, right), revealing an increased expression of *Pparγ* and *Retn*, and a trend towards increased expression of *Fabp4* and *Adipoq*, whereas *Lep* and *Nampt* expression remained unchanged. Collectively, these results indicate that eWAT of Yanomami-HMA mice contains more healthy, differentiated and less-inflamed lipid-storing adipocytes. Although no clear pathology could be detected in intestinal tissue of Belgian- and Yanomami-HMA mice (Suppl. Fig. 3f), expression of inflammatory markers *Ly6c*, *Il-1β* and *Il-6* was reduced in colon tissue of Yanomami-HMA mice (Suppl. Fig. 2c). Colonic expression of other inflammatory markers (*Ly6g*, *F4/80, Tnf*) was not significantly different.

Consistent with the reduced hepatic and adipose tissue pathology, Yanomami-HMA mice exhibited an improved systemic metabolic profile. Compared with Belgian-HMA mice, circulating free fatty acids (FFA; Fig. 2l) concentrations were significantly reduced, together with lower serum alanine aminotransferase activity (ALT, Fig. 2m) indicative of reduced hepatocellular injury. The metabolically favourable phenotype was accompanied by significantly lower circulating levels of inflammatory cytokines IL-6 and IL-17, and a trend towards decreased TNF in Yanomami-HMA mice (Fig. 2n), indicating reduced systemic inflammation. In contrast, no significant differences were detected in serum IFNγ or MCP-1 levels (Suppl. Fig. 2d) or serum levels of diabetes markers insulin, leptin, resistin and PAI-1 (Suppl. Fig. 2e). Interestingly, ghrelin levels were increased in serum of Yanomami-HMA mice. Of note, serum used in FFA/ALT/cytokine quantification was collected 120 minutes after IPGTT following an overnight fast, which makes these read-outs reflective of acute glucose/stress context rather than a basal HFD steady state. Metabolic hormones were measured in fasting serum to represent the basal HFD steady state.

### The Yanomami microbiota reprograms hepatic metabolic gene expression

Given the significant difference in liver weight and hepatic lipid accumulation between Belgian- and Yanomami-HMA mice, we used bulk mRNA sequencing to investigate whether these phenotypic changes were accompanied by transcriptional reprogramming of the liver. Principal component analysis (PCA) revealed a clear separation between the 2 groups (Fig. 2o). Differential expression analysis identified a total of 986 differentially expressed genes (DEG), indicating extensive microbiota-dependent remodelling of the hepatic transcriptome (Fig. 2p, top DEGs shown in Suppl. Fig. 2f). Among the most strongly downregulated genes in Yanomami-HMA mice were *Cidea* and *Cidec,* two key regulators of hepatic lipid droplet fusion and triglyceride storage^38^, consistent with the reduced accumulation of large lipid droplets observed histologically (Fig. 2h). Conversely, Yanomami-HMA livers exhibited marked upregulation of a cluster of major urinary protein (*Mup*) genes, alongside *Hsd3b5* and *Serpina1e* (Fig. 2p and Suppl. Fig. 2f). As hepatic *Mup* expression has been linked to improved glucose tolerance and insulin sensitivity^39^, these transcriptional changes are consistent with the metabolically favourable phenotype of Yanomami-HMA mice.

To identify the biological pathways associated with these transcriptional changes, we performed gene set enrichment analysis (GSEA). This analysis revealed an upregulation of pathways associated with oxidative phosphorylation, fatty acid metabolism, and adipogenesis in the Belgian group, and an upregulation of bile acid metabolism in the Yanomami group, among others (Fig. 2q). Consistent with these pathway-level changes, genes involved in lipid droplet formation and lipid uptake were significantly downregulated in the Yanomami group, including *Plin2 and Lpl*. These changes were accompanied by increased expression of *Ppargc1b*, a regulator of mitochondrial biogenesis and oxidative capacity^40^, and decreased expression of *Fgf21* (Suppl. Fig. 2g), a hepatokine previously suggested as potential biomarker for liver disease^41^, both consistent with reduced lipid storage and altered oxidative metabolism seen in the Yanomami-HMA mice. Two additional lipid-handling genes, *Fabp4* and *Trem2*, were significantly downregulated in the liver of Yanomami-HMA mice (Suppl. Fig. 2g), in line with previous gene expression data from eWAT (Fig. 2k and Suppl. Fig. 2b). Furthermore, Yanomami-HMA mice show increased hepatic expression of *Abcc2* and *Sult2a8*, both involved in bile acid metabolism^42,43^. These findings support a shift toward increased bile acid transport and detoxification capacity in the livers of Yanomami-vs. Belgian-HMA mice.

To identify upstream transcriptional regulators potentially driving these expression changes, we performed transcription factor activity prediction analysis (Suppl. Fig. 2h). This analysis identified increased activity of *Hnf1a* and *Foxa2* in livers of Yanomami-HMA mice, which regulate bile acid and lipid metabolic gene programs^44,45^, alongside decreased predicted activity of *Mlxipl*, a principal transcriptional driver of de novo lipogenesis^46^. This analysis supports the increased bile acid metabolism and decreased adipogenesis observed in livers of Yanomami-HMA mice.

### Cohort-level assessment of the protective effects of Yanomami microbiota

To determine whether the protective metabolic phenotype observed in the representative donor set generalized across Yanomami donors, we repeated the HFD experiment using additional Belgian and Yanomami donors, selected to achieve broadly comparable demographic characteristics, with similarity in sex, age and BMI status prioritized where possible (Table 1). Donors were treated as independent biological replicates in all statistical analyses. Donors were selected to represent the overall microbiota composition of their respective cohorts, excluding donor sets that occupied extreme ends of the diversity spectrum, ensuring that the subsequent HMA experiments evaluated microbiota characteristics representative of each human cohort (Fig. 1d).

Across donor-level pooled analysis, transplantation of Yanomami microbiota was associated with significantly different trajectories of HFD-induced weight gain compared with Belgian microbiota (Time × Group interaction, p = 0.011), with a smaller increase in body weight observed in the Yanomami group (Fig. 3a), demonstrating that the protective metabolic phenotype extends beyond a single representative donor set. As expected, the magnitude of protection varied among individual donor sets. Most Belgian microbiota promoted substantially greater weight gain than their Yanomami counterparts, whereas two Belgian donors (BEL_016 and BEL_039) induced relatively limited weight gain (Suppl. Fig. 3a, b). Similarly, random non-fasted blood glucose measurements varied considerably between donor sets (Suppl. Fig. 3c), while recipients of Yanomami microbiota exhibited a trend towards lower fasting glucose concentrations when donor sets were considered the biological unit of analysis (Fig. 3b). Classification according to established fasting glucose criteria for diabetes (>200 mg/dL^47^) revealed a significantly lower incidence of diabetes among recipients of Yanomami microbiota (Fig. 3c). When looking at separate donor sets, we see lower fasting glucose concentrations in three out of seven donor sets following overnight fasting (Fig. 3d). Likewise, liver weight was reduced in five out of seven donor sets (Fig. 3f), although this effect did not reach statistical significance following donor-level pooling (Fig. 3e), whereas eWAT weight remained comparable between groups (Fig. 3g).

**Fig. 3.**
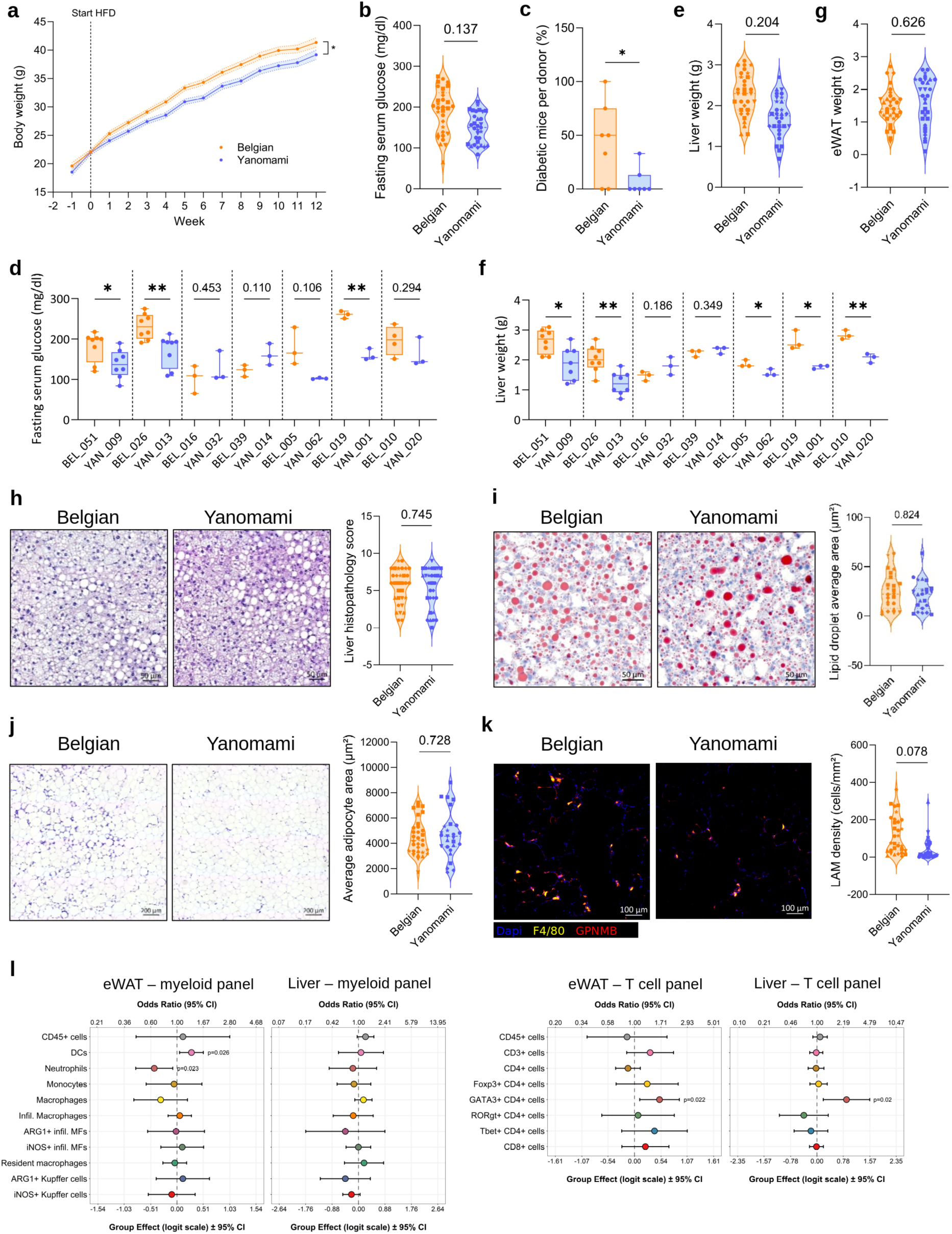
Yanomami gut microbiota reduces body weight gain and diabetes development following HFD at the cohort level. **a,** Weight follow-up over the course of high-fat diet (HFD) administration. Data are represented as mean ± SEM. Data were analysed using a linear mixed-effects model fitted by restricted maximum likelihood (REML) with fixed effects for Experiment, Time, Group, and Time x Group interaction, corrected for start weight, and random effects for mouse ID (repeated measures), donor, and donor–time interaction. Indicated significances correspond to the Time × Group interaction (P = 0.011; group effect: P = 0.07). **b,e,g,** Significance was determined using a linear mixed-effects model fitted by REML (details in methods section, P values indicated). **b,** Fasting serum glucose measurements following 12 weeks of HFD. Each datapoint represents a mouse, each symbol represents a different Belgian or Yanomami human microbiota. Mice with the same microbiota are treated as technical replicates. **c,** Frequency of diabetic mice within each donor. The cutoff for diabetes development was set at a fasting serum glucose level of >= 200mg/dl. Significance was determined using a Mann-Whitney test (Hodges-Lehmann difference = −37.00; Mann-Whitney U = 9.5, n = 7 per group, exact two-tailed P = 0.0495). **d,** Fasting serum glucose levels of separate donor sets following 12 weeks of HFD. Significant differences were determined between donors using a Welch’s t test (P values indicated). **e,** Liver weight measured following 12 weeks of HFD. Each datapoint represents a mouse, each symbol represents a different Belgian or Yanomami human microbiota. Mice with the same microbiota are treated as technical replicates. **f,** Liver weight of separate donor sets following 12 weeks of HFD. Significant differences were determined between donors using a Welch’s t test (P values indicated). **g,** eWAT weight measured following 12 weeks of HFD. Each datapoint represents a mouse, each symbol represents a different Belgian or Yanomami human microbiota. Mice with the same microbiota are treated as technical replicates. **h,** Representative liver histology images (H&E, left) of Belgian vs. Yanomami mice following 12 weeks of HFD, with associated liver pathological scoring (right). Each datapoint represents a mouse, each symbol represents a different Belgian or Yanomami human microbiota. Mice with the same microbiota are treated as technical replicates. **i,** Representative liver histology images (Oil Red O, left) of Belgian vs. Yanomami mice following 12 weeks of HFD, with associated measurement of lipid droplet area (stained red on image, right). Each datapoint represents the average of measured lipid droplets in 3 regions of a liver section of one mouse, each symbol represents a different Belgian or Yanomami human microbiota. Mice with the same microbiota are treated as technical replicates. **j,** Representative eWAT histology images (H&E, left) of Belgian vs. Yanomami mice following 12 weeks of HFD, with associated measurement of adipocyte area (right). Each datapoint represents the average of measured adipocytes in an eWAT section of one mouse, each symbol represents a different Belgian or Yanomami human microbiota. Mice with the same microbiota are treated as technical replicates. **k,** Representative eWAT immunofluorescence images (blue: Dapi, yellow: F4/80, red: GPNMB) of Belgian vs. Yanomami mice following 12 weeks of HFD, with associated measurement of LAM density (F4/80+ GPNMB+ cells, right). Each datapoint represents the average of measured LAM density in 3 regions of an eWAT section of one mouse, each symbol represents a different Belgian or Yanomami human microbiota. Mice with the same microbiota are treated as technical replicates. **l,** Flow cytometry analysis of innate (left) and adaptive (right) immune compartments in the eWAT and liver of mice with a Yanomami vs. Belgian microbiota. Analysis was performed on 5 donor sets for the innate eWAT/liver and adaptive liver panel, and for 4 donor sets for the adaptive eWAT panel. Dots represent differences (effect sizes) in immune cell proportions between Yanomami and Belgian mice (represented by vertical line) shown on the logit scale (log odds ratio; lower X-axis) and on the odds ratio (OR) scale (higher X-axis). Colours denote the various immune cell types. Horizontal lines show 95% confidence intervals. The vertical line indicates no effect (OR = 1). MF: macrophage. *: P < 0.05

We next assessed glucose tolerance by intraperitoneal glucose tolerance testing (IPGTT) for each donor set individually (Suppl. Fig. 3c). Recipients of Yanomami microbiota exhibited better glucose tolerance, reflected by a reduced area under the curve (AUC), in four out of seven donor sets, reaching statistical significance for the BEL_019–YAN_001 donor set. Conversely, two Belgian microbiota (BEL_016 and BEL_039) showed faster glucose metabolism than their Yanomami counterparts, consistent with their comparatively limited weight gain, although this only becomes significant for the BEL_016–YAN_032 donor set. Together, these findings indicate that while the magnitude of improvement in glucose tolerance varies among individual microbiota, the observed donor heterogeneity closely mirrors the variability in HFD-induced weight gain.

Histological analysis of liver tissue from pooled experiments confirms the presence of hepatocellular hypertrophy with micro- and macrovesicular steatosis in both Belgian- and Yanomami-HMA mice, as before, with no differences between groups (Fig. 3h). Analysis of lipid droplet area of all pooled samples does not reveal a difference in lipid droplet size in the liver of Belgian-vs. Yanomami-HMA mice (Fig. 3i). In addition, there is no difference in average adipocyte area (Fig. 3j) and the quantification of LAM density reveals a trend towards a decrease in LAM infiltration in the eWAT of Yanomami mice, although this does not reach significance (Fig. 3k). Finally, intestinal histology was compared, whereby neither colon nor ileum histology showed clear differences (Suppl. Fig. 3d), nor did we detect any differences in colonic mucus thickness (measured on Carnoy-fixed, AB/PAS-stained sections, Suppl. Fig. 3e).

To determine whether the metabolic differences between Belgian- and Yanomami-HMA mice were accompanied by alterations in tissue immunity, we performed immunophenotyping of liver and eWAT from five independent donor set experiments (Fig. 3l). This includes an analysis of innate (myeloid) and adaptive (T cell) immunity (expressed as odds ratios of frequencies, gating strategy shown in Suppl. Fig. 4). While total leukocyte (CD45+ cells) infiltration was comparable between groups, Yanomami-HMA mice exhibited a distinct remodelling of adipose immunity, characterized by altered innate and adaptive immune compartments. We observed reduced neutrophil and increased dendritic cell frequencies within eWAT (Fig. 3l, left), which was not observed in the liver. Given the established role of macrophages in obesity-associated metabolic dysfunction^48^, we characterized the macrophage population in detail in both eWAT and liver. Neither total macrophages (MF), nor infiltrating (Vsig4-) or resident (Vsig4+) macrophages showed any differences in eWAT or liver of Belgian- and Yanomami-HMA mice. Furthermore, expression of polarization markers iNOS and ARG1 was comparable in both groups in liver and eWAT. Interestingly, analysis of adaptive immunity revealed a consistent increase in GATA3⁺ CD4⁺ T cells in both liver and eWAT of Yanomami-HMA mice (Fig. 3l, right), indicating a consistent shift towards type 2 immunity. In contrast, CD8⁺ T cells, Th1 (T-bet⁺), Th17 (RORγt⁺) and regulatory T cells (Foxp3⁺) remained unchanged. These findings demonstrate that transplantation of Yanomami microbiota induces selective remodelling of innate immune cell populations within adipose tissue together with a systemic Th2-biased adaptive immune response.

Analysis of circulating metabolic and inflammatory biomarkers revealed limited differences between Belgian- and Yanomami-HMA mice. Serum concentrations of insulin, leptin, resistin, ghrelin and PAI-1 were comparable between groups, although insulin levels tended to be lower in recipients of Yanomami microbiota (Suppl. Fig. 5a). Likewise, serum ALT activity showed a trend towards lower levels in Yanomami-HMA mice without reaching statistical significance (Suppl. Fig. 5b). Serum FFA concentrations and pro-inflammatory cytokine levels were also comparable between groups (Suppl. Fig. 5c,d). Since immune profiling revealed a Th2 skewing in both eWAT and liver tissue, we measured serum IL-13 levels to confirm systemic presence of Th2 cytokines. However, these measurements failed to reach the detection limit and were therefore excluded.

Finally, we evaluated tissue-specific gene expression by RT-qPCR using pooled RNA from individual donor sets. In contrast to the detailed analysis of the representative BEL_026–YAN_013 donor set (Fig. 2), donor-level pooled analysis did not reveal significant differences in expression of genes associated with adipose tissue metabolism, adipokine production or inflammation in eWAT (Suppl. Fig. 5e), nor in genes related to intestinal barrier function, gut hormones or inflammation in the colon (Suppl. Fig. 5f).

Collectively, these validation experiments demonstrate that transplantation of Yanomami microbiota significantly reduces HFD-induced body weight gain across seven independent donor sets, establishing protection against diet-induced weight gain as a cohort-level phenotype. Although expected inter-individual donor variation, particularly among Belgian microbiota, attenuated the statistical significance of several secondary metabolic and molecular readouts following donor-level pooling, the overall direction of effect remained remarkably consistent across multiple independent endpoints and was accompanied by reproducible alterations in adipose tissue immune composition and Th2 skewing.

### The Yanomami-derived microbiota exhibits increased ecological stability following HFD challenge

To investigate the ecological mechanisms underlying the phenotype of the representative donor set (Fig. 2, BEL_026–YAN_013), we profiled faecal metagenomes in recipient mice before and after HFD challenge. We found that HFD challenge induced markedly different ecological responses depending on donor microbiota origin. PCoA demonstrated that community composition was strongly influenced by both donor origin and diet (Fig. 4a). Although significant compositional shifts occurred in both groups following HFD, paired analysis revealed substantially greater ecological restructuring of Belgian-than Yanomami-derived communities (Fig. 4b), indicating that the Yanomami microbiota remained considerably more stable following dietary perturbation. Consistent with this increased ecological stability, alpha diversity responded differently to HFD according to donor microbiota origin. Shannon and Gini-Simpson diversity increased significantly in Belgian-HMA mice following HFD, whereas neither diversity metric changed in Yanomami-HMA mice (Fig. 4c). Consequently, Shannon diversity was comparable between Belgian- and Yanomami-derived communities prior to HFD but became significantly higher in Belgian-HMA mice after dietary intervention.

**Fig. 4.**
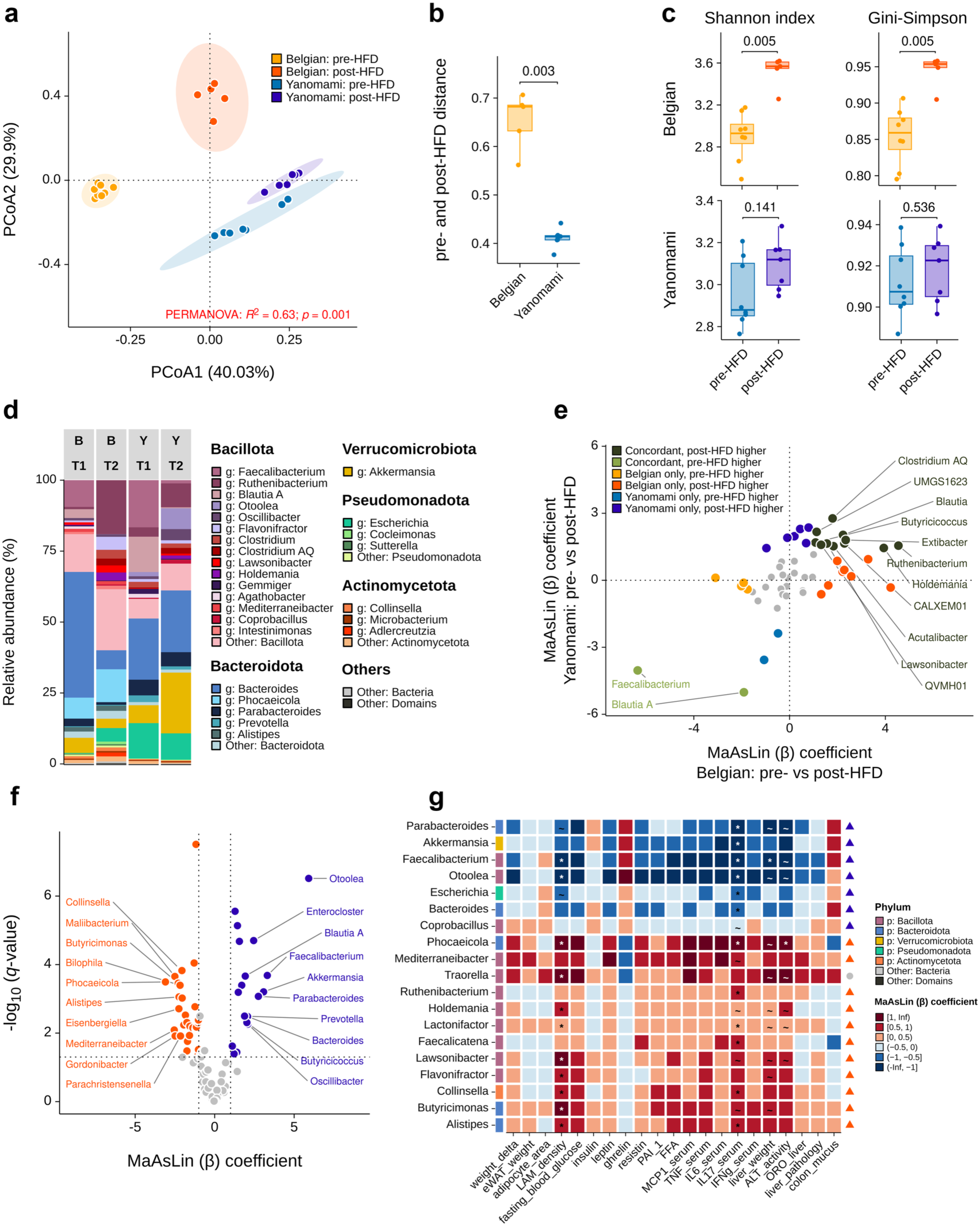
Belgian- and Yanomami-derived gut microbiota (BEL_026 vs. YAN_013) differ in taxonomic composition and respond differently to high-fat diet. **a,** Principal coordinate analysis (PCoA) of Bray–Curtis dissimilarities calculated from unaggregated taxonomic features. Community structure was associated with donor cohort and time point (PERMANOVA: R^2^ = 0.632, F = 21.48, P = 0.001). Pairwise differences were detected between all groups: Belgian pre- versus post-HFD (R^2^ = 0.682, P = 0.001), Belgian versus Yanomami pre-HFD (R^2^ = 0.677, P = 0.002), Belgian pre-HFD versus Yanomami post-HFD (R^2^ = 0.818, P = 0.001), Belgian post-HFD versus Yanomami pre-HFD (R^2^ = 0.636, P = 0.002), Belgian versus Yanomami post-HFD (R^2^ = 0.654, P = 0.002), and Yanomami pre- versus post-HFD (R^2^ = 0.524, P = 0.001). Within-group dispersion also differed (P < 0.001), indicating that both group centroids and dispersion may contribute to these differences. **b,** Bray–Curtis dissimilarity between paired pre- and post-HFD samples from mice with Belgian (n = 5) or Yanomami (n = 7) microbiota. Yanomami-derived communities showed a smaller compositional shift (Wilcoxon rank-sum test, P = 0.0025, estimated difference = 0.265, 95% CI 0.155–0.292). **c,** Genus-level Shannon and Gini–Simpson diversity before and after high-fat diet (HFD) administration in mice colonized with Belgian (top; pre-HFD, n = 8; post-HFD, n = 5) or Yanomami microbiota (bottom; pre-HFD, n = 8; post-HFD, n = 7). Diversity increased after HFD in Belgian-derived communities (Shannon: FDR-adjusted P = 0.0047, estimated pre- versus post-HFD difference = −0.640, 95% CI −0.904 to −0.373; Gini–Simpson: FDR-adjusted P = 0.0047, difference = −0.0851, 95% CI −0.151 to −0.0501), but not in Yanomami-derived communities (Shannon: adjusted P = 0.141; Gini–Simpson: adjusted P = 0.536; Wilcoxon rank-sum tests).**d,** Mean genus-level relative abundance before (T1) and after (T2) HFD. The 15 most abundant Bacillota genera, five most abundant Bacteroidota genera, and three most abundant genera from each of Verrucomicrobiota, Pseudomonadota, and Actinomycetota are shown. Remaining bacterial taxa were grouped as “Other: Bacteria” and non-bacterial taxa as “Other: Domains”. **e,** Differential-abundance responses to HFD among genera detected in both donor groups. Genera associated with the same time point in both groups are shown in shades of green; grey points indicate genera without significant associations. **f,** Differentially abundant genera between Yanomami- and Belgian-derived communities after HFD (Belgian, n = 5; Yanomami, n = 7). Positive and negative MaAsLin2 coefficients (β) indicate associations with Yanomami- and Belgian-derived communities, respectively. In **e–f**, significance was defined as |β| > 1 and BH-adjusted q < 0.05; the ten strongest positive and negative associations are labelled in **f**. **g,** Associations between genera and host metabolic, inflammatory, and histopathological measurements after HFD. Cell colours indicate MaAsLin2 coefficient direction and magnitude; symbols indicate BH-adjusted significance levels (∼, q < 0.01; *, q < 0.05). The left annotation indicates phylum, while the right annotation indicates whether each taxon was associated with Belgian- or Yanomami-derived microbiota according to the differential abundance analysis. Y-axes were ordered by hierarchical clustering using Euclidean distance and complete linkage to improve visualization. eWAT, epididymal white adipose tissue; FFA, free fatty acids; IL, interleukin; IFN-γ, interferon-γ; MCP-1, monocyte chemoattractant protein 1; PAI-1, plasminogen activator inhibitor 1; TNF, tumour necrosis factor.

Recipient microbiota diverged from the original donor stool and gavage inoculum following engraftment, yet remained highly similar among mice colonized with the same donor microbiota (Suppl. Fig. 6a). Interestingly, despite exhibiting greater overall ecological stability, Yanomami-derived communities retained a smaller fraction of donor-core genera following HFD than Belgian-derived communities (40.4% versus 76.0%; Suppl. Fig. 6b: square icons), indicating that community restructuring and donor-core retention reflect distinct ecological properties.

Despite common responses to HFD, Belgian- and Yanomami-derived microbiota adopted distinct community configurations following dietary intervention (Fig. 4d–f). Several abundant genera such as *Ruthenibacterium*, *Clostridium AQ*, *Lawsonibacter*, and *Holdemania* increased after HFD in both groups, whereas *Blautia A* and *Faecalibacterium* decreased, indicating conserved microbial responses to HFD. However, the magnitude and resulting composition of these shifts differed substantially according to microbiota origin. Belgian-HMA communities showed a marked reduction in *Bacteroides* and *Faecalibacterium*, together with an expansion of *Ruthenibacterium* post-HFD (Fig. 4d) and enrichment of genera including *Phocaeicola*, *Mediterraneibacter*, *Alistipes*, *Bilophila* and *Collinsella* compared to Yanomami-HMA communities post-HFD (Fig. 4f). In contrast, Yanomami-HMA communities retained higher *Bacteroides* abundance and showed increased *Akkermansia* upon HFD (Fig. 4d), with post-HFD enrichment of *Otoolea*, *Parabacteroides*, *Oscillibacter*, *Prevotella* and *Faecalibacterium* in relation to Belgian-microbiota recipients (Fig. 4f).

To explore whether these cohort-specific taxa were associated with host physiology, we correlated bacterial abundances with metabolic and inflammatory parameters (Fig. 4g). Notably, several genera enriched in Yanomami-derived microbiota following HFD, including *Otoolea*, *Faecalibacterium*, *Akkermansia* and *Parabacteroides* were negatively associated with serum IL-17 concentrations. Furthermore, *Otoolea*, *Faecalibacterium* and *Parabacteroides* were negatively associated with ALT activity. *Faecalibacterium* and *Otoolea* also correlated negatively with LAM density and *Faecalibacterium* with relative liver weight. In contrast, the Belgian-associated genus *Phocaeicola* was positively associated with all four parameters. Together, these analyses suggest *Otoolea*, *Faecalibacterium* and *Parabacteroides* as candidate taxa associated with the metabolically protective phenotype conferred by Yanomami-derived microbiota.

To determine whether these ecological responses extended beyond the representative donor set (BEL_026 vs. YAN_013), metagenomic profiles were summarised to one profile per human donor across all HMA experiments. Belgian-derived communities again retained a larger fraction of donor-core genera following HFD than Yanomami-derived communities (Suppl. Fig. 6b). Likewise, Shannon and Gini– Simpson diversity increased significantly following HFD in Belgian-derived microbiota, whereas neither diversity metric changed in Yanomami-derived communities (Suppl. Fig. 6c). Community composition remained significantly associated with donor origin and dietary intervention (Suppl. Fig. 6d). Although the greater pre-to post-HFD compositional shift of Belgian-derived microbiota no longer reached statistical significance after donor-level summarization (Suppl. Fig. 6e), the overall direction of effect remained consistent with increased ecological stability of Yanomami-derived microbiota.

Differential-abundance analyses likewise identified both conserved and donor-dependent microbial responses to HFD. Consistent with the representative donor experiment, *Ruthenibacterium* increased whereas *Faecalibacterium* decreased following HFD in both Belgian and Yanomami groups, while several population-specific taxa remained associated with Belgian- or Yanomami-derived microbiota. Belgian-derived post-HFD profiles were enriched in genera including *Eisenbergiella*, *Adlercreutzia* and *Mediterraneibacter*, whereas *Hominicoprocola*, *Odoribacter*, *Butyricicoccus*, *Bulleidia* and *Prevotella* were associated with Yanomami-derived profiles (Suppl. Fig. 6f,g). Despite expected variation in the taxa responding within individual donor sets, the most discriminating genera continued to separate donor-level profiles according to microbiota origin and dietary status (Suppl. Fig. 6h). By contrast, correlation analysis with donor-level host metabolic parameters and immune populations did not identify robust associations after donor-level averaging (Suppl. Fig. 6i,j), reflecting the increased biological heterogeneity across independent human donors. These findings identify increased ecological stability as a reproducible cohort-level characteristic of Yanomami-derived microbiota following HFD challenge, providing a potential ecological basis for their reduced susceptibility to diet-induced metabolic dysfunction.

### Yanomami-derived microbiota exhibit greater functional resilience and distinct metabolic responses to high-fat diet

To determine whether the greater taxonomic resilience observed in the representative BEL_026 vs. YAN_013 donor set was accompanied by specific changes in microbial functional potential, we compared predicted functional profiles before and after HFD challenge. Bray–Curtis analysis of KEGG ortholog (KO) profiles revealed a strong effect of both microbiota origin and dietary status (Fig. 5a). As observed for taxonomic composition, within-group dispersion also differed, indicating some contribution of intra-group variation. Paired analysis demonstrated that Belgian-derived microbiota underwent a significantly larger functional shift following HFD than Yanomami-derived microbiota (Fig. 5b). The greater taxonomic stability of Yanomami-derived communities was accompanied by preservation of their predicted functional repertoire during HFD-challenge. This difference was further reflected in the extent of HFD-associated functional reprogramming. Functional divergence between Belgian- and Yanomami-derived communities became even more pronounced following HFD, with 2,544 KOs associated with Belgian-derived communities compared with 1,054 associated with Yanomami-derived communities (Fig. 5c). Belgian-derived communities contained 1,120 KOs associated with the pre-HFD state and 2,868 KOs associated with the post-HFD state, whereas Yanomami-derived communities contained only 830 and 781 pre- and post-HFD associated KOs, respectively (Fig. 5d,e). Differentially abundant KOs were distributed across multiple KEGG metabolic subclasses, with carbohydrate metabolism representing the largest category in both groups but showing substantially broader representation among Belgian-derived communities (Fig. 5f). Together, these results indicate that the greater taxonomical restructuring of the Belgian-derived microbiota is accompanied by more extensive remodelling of their encoded metabolic repertoire.

**Fig. 5.**
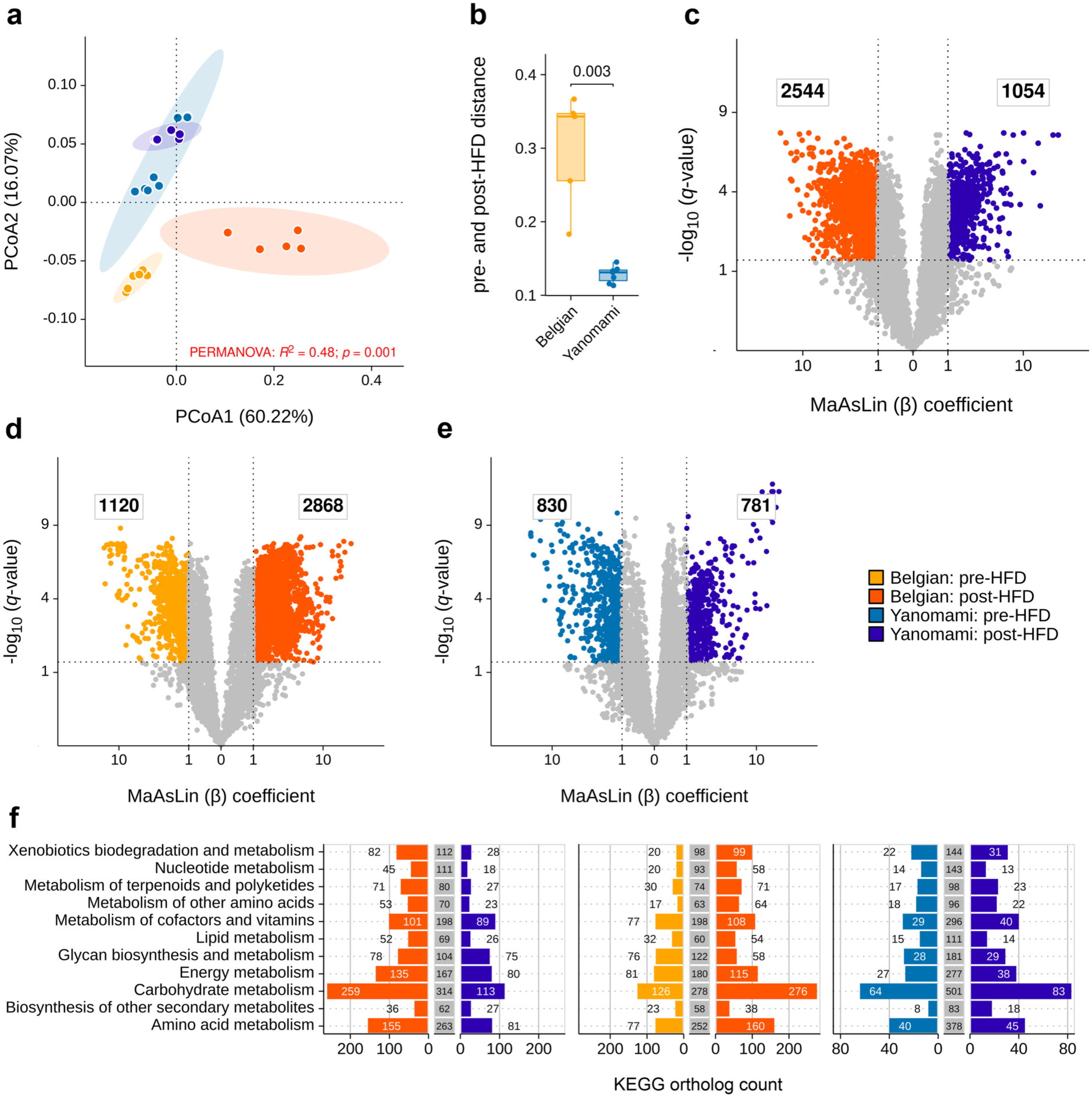
Belgian- and Yanomami-derived gut microbiota (BEL_026 vs. YAN_013) show distinct functional potential following high-fat diet. **a**, PCoA of Bray–Curtis dissimilarities calculated from KEGG ortholog (KO) profiles (Belgian: pre-HFD, n = 8; post-HFD, n = 5; Yanomami: pre-HFD, n = 8; post-HFD, n = 7). Functional composition was associated with donor cohort and time point (PERMANOVA: R^2^ = 0.480, F = 11.52, P = 0.001). FDR-corrected pairwise differences were detected between all groups: Belgian pre- versus post-HFD (R^2^ = 0.823, adjusted P = 0.001), Belgian versus Yanomami pre-HFD (R^2^ = 0.498, adjusted P = 0.001), Belgian pre-HFD versus Yanomami post-HFD (R^2^ = 0.780, adjusted P = 0.001), Belgian post-HFD versus Yanomami pre-HFD (R^2^ = 0.672, adjusted P = 0.001), Belgian versus Yanomami post-HFD (R^2^ = 0.778, adjusted P = 0.003), and Yanomami pre- versus post-HFD (R^2^ = 0.389, adjusted P = 0.001). Within-group dispersion also differed (P < 0.001), indicating that both group centroids and dispersion may contribute to these differences. **b**, Bray–Curtis dissimilarity between paired pre- and post-HFD KO profiles from mice with Belgian (n = 5) or Yanomami (n = 7) microbiota. Belgian-derived communities showed a greater functional shift (Wilcoxon rank-sum test, adjusted P = 0.0025, estimated difference = 0.210, 95% CI 0.0674–0.234). **c**, Differentially abundant KOs between Yanomami- and Belgian-derived communities after HFD. Positive and negative MaAsLin2 coefficients (β) indicate associations with Yanomami- and Belgian-derived communities, respectively. Significance was defined as |β| > 1 and BH-adjusted q < 0.05; 2,544 KOs were associated with Belgian-derived communities and 1,054 with Yanomami-derived communities. **d–e**, Differentially abundant KOs between pre- and post-HFD samples within Belgian-derived (**d**) and Yanomami-derived (**e**) communities. In Belgian-derived communities, 1,120 KOs were associated with pre-HFD and 2,868 with post-HFD samples; corresponding values for Yanomami-derived communities were 830 and 781, respectively. In c–e, significance was defined as |MaAsLin2 coefficient (β)| > 1 and BH-adjusted q < 0.05, and square-root-scaled axes were used for visualization. **f**, Distribution of differentially abundant KO assignments across KEGG metabolism subclasses for the post-HFD comparison between Belgian- and Yanomami-derived communities (left), the Belgian pre- versus post-HFD comparison (centre), and the Yanomami pre- versus post-HFD comparison (right). Labels indicate the number of differentially abundant KO assignments associated with each cohort or time point; central grey bars represent non-differentially abundant KOs.

Pathway-level analyses further highlighted distinct metabolic configurations following HFD. Belgian-derived communities were enriched in pathways involved in carbohydrate catabolism and sugar metabolism, including glycolysis/gluconeogenesis, pentose phosphate pathway, fructose and mannose metabolism, starch and sucrose metabolism, and pentose and glucuronate interconversions (Suppl. Fig. 7). In contrast, Yanomami-derived communities were enriched for pathways associated with central carbon metabolism, including tricarboxylic acid cycle, propanoate metabolism, pyruvate metabolism, and glyoxylate and dicarboxylate metabolism (Suppl. Fig. 7). Lipid metabolism displayed a similarly divergent pattern. Glycerolipid metabolism and secondary bile acid biosynthesis were associated with Belgian-derived communities, whereas sphingolipid metabolism and fatty acid biosynthesis were associated with Yanomami-derived communities (Suppl. Fig. 7). Additional differences involved amino acid metabolism, xenobiotic degradation, cofactor and vitamin metabolism, energy metabolism, and terpenoid and polyketide pathways (Suppl. Fig. 7). In particular, Yanomami-derived communities showed broader representation of pathways related to vitamin and cofactor biosynthesis, while Belgian-derived communities were associated with several xenobiotic degradation pathways. The taxonomic contributors underlying these functional differences also markedly differed between microbiota origins. In Yanomami-derived communities, predicted pathway abundance was primarily attributed to *Akkermansia*, *Bacteroides,* and *Parabacteroides*. In contrast, pathway abundance in Belgian-derived communities was distributed across a broader range of Bacillota taxa, including numerous low-abundance taxa or unclassified genera (grouped as “Other: Bacillota”; Suppl. Fig. 7), with HFD-associated taxa such as *Ruthenibacterium*, *Lawsonibacter* and *Holdemania* representing prominent contributions. *Escherichia* contributed consistently to pathways in both cohorts. Together, these findings indicate that the greater taxonomic remodelling of Belgian-derived communities following HFD was driven by a broader and compositionally distinct microbial community.

Across all donor sets, Belgian-derived communities showed a larger HFD-induced functional shift than Yanomami-derived communities (Suppl. Fig. 8a,b), confirming the pattern observed in the representative donor set, although the separation was less pronounced due to donor-to-donor variability. Consistent with this, functional composition differed significantly between Belgian pre- and post-HFD profiles, whereas no significant pre- to post-HFD separation was detected among Yanomami-derived communities (Suppl. Fig. 8a). By contrast, the relative distribution of broad KEGG metabolic subclasses remained comparatively similar between Belgian- and Yanomami-derived communities before and after HFD (Suppl. Fig. 8c). At the KO level, Belgian-derived profiles showed a substantially stronger HFD-associated response, with 674 KOs associated with pre-HFD and 1,102 with post-HFD profiles, compared with 43 and 118 KOs, respectively, in Yanomami-derived communities (Suppl. Fig. 8d,e). Thus, although Yanomami-derived microbiota were not completely functionally invariant, their HFD-associated KO-level response was markedly more limited than that observed in Belgian-derived communities. Pathway-level profiles nevertheless showed substantial inter-donor variation, particularly among Yanomami-derived microbiota, indicating that reduced aggregate functional change does not imply uniform responses across individual donors (Suppl. Fig. 8f). Overall, the donor-level analysis reproduced the stronger directional functional response of Belgian-derived communities and supports greater predicted functional stability of Yanomami-derived microbiota following HFD challenge.

## Discussion

Industrialization is associated with the progressive loss of microbial diversity and bacterial taxa that are common in non-urbanized populations implementing traditional subsistence practices^2,12,13^. Although these compositional changes have been extensively documented, whether they causally influence susceptibility to immune and metabolic disorders prevalent in industrialized societies, including obesity and type 2 diabetes, remains largely unknown. Here, we addressed this question by comparing gut microbiota from the Brazilian Yanomami, representing a low-industrialization Indigenous cohort maintaining a traditional lifestyle, with those from a Belgian urban-industrialized cohort, and investigated their impact on HFD-induced metabolic disease following transplantation into GF mice. This comparison was designed to contrast microbiota shaped by markedly different ecological and lifestyle contexts rather than to isolate the individual contributions of diet, lifestyle, geography or host genetics, all of which may influence microbiota composition.

In line with previous studies^13,49^, the Yanomami gut microbiota was overall more diverse than the Belgian microbiota and compositionally distinct, with enrichment of established VANISH taxa and depletion of BloSSUM taxa, supporting industrialization as a major determinant of gut microbiota composition across geographically distinct populations. Complementary assembly-based profiling showed that diversity differed among Yanomami populations, whereas taxonomic composition consistently distinguished each population from the Belgian cohort. This suggests that compositional remodelling more broadly characterises the contrast between Belgians and Yanomami than increased diversity alone (de Carvalho et al., *in prep*). Moreover, several Yanomami-enriched genera, including *Petrimonas, Duodenibacillus* and *Parolsenella*, have not previously been classified as VANISH taxa. Although validation across additional populations is required, our findings identify novel candidate low-industrialization-associated (VANISH) taxa and suggest that the microbial diversity depleted during industrialization remains incompletely catalogued.

Our findings demonstrate that Yanomami-derived microbiota reduce susceptibility to diet-induced weight gain and metabolic dysfunction, although the magnitude of these effects varies across individual donors. This protection was accompanied by a distinct microbial response to dietary perturbation. In the reference donor set, Yanomami derived microbial communities underwent smaller taxonomic and functional shifts following HFD than Belgian-derived microbial communities, while across all donors the reduced functional shift remained significant and the taxonomic response showed the same overall direction. These findings suggest that greater stability of microbial community function during nutritional perturbation may represent an important property of Yanomami-derived microbiota and raise the possibility that preservation of microbial functions contributes to improved host adaptation to HFD.

Detailed analysis of the representative donor set indicated altered adipose-liver lipid handling. Yanomami-HMA mice showed lower fasting serum glucose and liver weight, fewer crown-like structures in eWAT, and lower eWAT *Trem2* and *Tnf* expression, consistent with TREM2’s established role in conferring lipid-handling capacity to LAMs and driving their recruitment around damaged adipocytes^50^. Yanomami-HMA eWAT also showed a trend towards higher relative expression of adipogenic/lipid-storage genes (*Pparγ, Retn, Fabp4, Adipoq*). These findings are compatible with the adipose tissue expandability hypothesis^51^, whereby more effective lipid storage in adipose tissue limits lipid spillover to ectopic organs like the liver. Consistent with this interpretation, Belgian-HMA mice have higher circulating free fatty acids and increased hepatic expression of lipid-storage genes including *Cidea, Cidec, Plin2* and *Lpl*, and elevated serum ALT. Together, these findings suggest a link between impaired lipid-buffering, increased systemic spillover and hepatic lipid accumulation in Belgian-HMA relative to Yanomami-HMA mice, although this mechanistic link remains speculative and warrants direct testing.

The considerable inter-individual variability observed throughout this study highlights the complexity of host-microbiota interactions. Although Yanomami-derived microbiota conferred protection at the cohort level, individual donor communities varied substantially in both their metabolic effects and ecological responses to high-fat diet. Notably, Belgian-HMA mice derived from donors BEL_016 and BEL_039 clustered more closely with the Yanomami phenotype than with the remaining Belgian donors, emphasizing that host responses are determined by the composition and functional capacity of individual microbial communities rather than cohort origin per se. Interestingly, inter-individual variation appeared greater within the Belgian cohort than among the Yanomami donors, consistent with the broader dietary, environmental and lifestyle heterogeneity that characterizes industrialized populations. These observations further suggest that metabolically favourable host responses may arise through multiple microbial community configurations. Distinct bacterial taxa and functional repertoires may therefore converge on overlapping physiological outputs, reflecting a degree of functional redundancy within the gut ecosystem. This variability highlights the importance of identifying the microbial functions and molecular mechanisms that ultimately determine host physiology.

Beyond their metabolic effects, Yanomami-derived microbiota also induced an immunological phenotype characterized by an increased frequency of GATA3⁺ CD4⁺ T cells in both eWAT and liver under HFD. This suggests that modulation of type 2 immunity may represent one mechanism through which these microbiota influence host physiology. Adipose-resident CD4+ T cells have been implicated in metabolic homeostasis during diet-induced obesity, with a shift from a Th2-towards a Th1-dominated immunity associated with insulin resistance^52^. GATA3+ CD4+ T cell enrichment observed in Yanomami-HMA mice is therefore consistent with their favourable metabolic phenotype, although further research is needed to establish causality. Its reproducibility across multiple donor sets nevertheless suggests that modulation of type 2 immunity may represent a conserved functional property of Yanomami-derived microbiota.

A major next step will be to identify the specific microorganisms and microbial functions responsible for the observed phenotypes. The associations with individual taxa and predicted metabolic pathways identified here provide important candidates but do not establish causality. Mechanistic validation will require reductionist gnotobiotic approaches, including mono-colonization experiments and the construction of defined minimal microbial consortia in which candidate organisms can be systematically added, removed or genetically modified. Achieving this will depend on establishing comprehensive culture collections of bacteria isolated from Yanomami donors, providing the experimental resource needed to dissect the contributions of individual species and microbial communities to host physiology. Such approaches will determine whether protection is mediated by individual organisms, cooperative community interactions or functionally redundant taxa occupying similar ecological niches.

Equally important, the metagenomic analyses performed here define the metabolic repertoire encoded by the transplanted communities rather than their physiological activity. Integrated metabolomic and metatranscriptomic profiling will therefore be essential to identify transcriptionally active pathways and the microbial molecules that reach the intestinal lumen, circulation and host tissues. Combining these measurements with host transcriptomics, spatial technologies and immune profiling will enable direct reconstruction of the microbial-metabolic–host pathways underlying protection, with particular attention to established immunometabolic mediators such as SCFAs, bile acid derivatives and tryptophan metabolites, as well as to previously uncharacterized microbial molecules and biosynthetic pathways uniquely preserved in low-industrialization microbiota. Notably, SCFA- and bile-acid-related functions contributed to the predicted functional differences involving Yanomami HMA communities and were also altered in the donor source populations, indicating that these signals emerge at the population level and remain detectable after transplantation (de Carvalho et al., *in prep*). Identifying these conserved protective functions and their underlying mechanisms may ultimately provide opportunities for microbiota-inspired therapeutic strategies, highlighting low-industrialization microbiota as a valuable reservoir of biological functions that have become rare or potentially lost during industrialization.

Although Yanomami-HMA mice exhibited reduced weight gain, the relative contributions of food intake, nutrient absorption, physical activity and energy expenditure could not be resolved because metabolic-cage studies were not feasible within our gnotobiotic facility. This may be particularly relevant to the unexpectedly higher serum ghrelin — the main orexigenic hormone, which stimulates eating and promotes fat storage^53^ — observed in Yanomami-vs. Belgian-HMA mice, a finding seemingly at odds with the lower weight gain of Yanomami-HMA mice. We speculate that this increased ghrelin could reflect a compensatory response to altered energy absorption or expenditure. Future experiments incorporating indirect calorimetry, locomotor activity, food intake and faecal energy measurements will therefore be important to distinguish reduced energy acquisition from increased energy expenditure or altered substrate utilization, and to test this hypothesis directly.

It is important to note that research involving Indigenous microbiota must be conducted through respectful, equitable and sustained partnerships with the Indigenous communities participating in the research^54^. This requires compliance with all relevant ethical, legal and access-and-benefit-sharing frameworks, transparent governance of biological samples and data, and continuous dialogue with local communities throughout the research process. Scientific objectives should never supersede Indigenous rights, autonomy or cultural values. Supporting Indigenous communities in maintaining cultural continuity, territorial sovereignty, and self-determined ways of life is intrinsically important. Separately, understanding how industrialization affects microbial diversity may help characterize the ecological consequences of rapid social and environmental change.

Together, our findings demonstrate that microbiota from a low-industrialization cohort harbour ecological and metabolic properties that improve host adaptation to nutritional stress. Yanomami-derived microbiota were associated with reduced weight gain and diabetes incidence across donor sets, albeit with substantial inter-donor variability, and these outcomes corresponded to greater ecological and functional resilience, including enrichment for SCFA-oriented taxa and metabolism, relative to the more extensively remodelled Belgian-derived communities. By linking microbial ecological resilience with protection against diet-induced metabolic dysfunction, this study provides a framework for identifying microbial functions lost during industrialization. Elucidating the responsible organisms, community interactions and molecular mediators may ultimately enable the development of microbiota-inspired therapeutics while underscoring the importance of conducting such research through equitable, community-engaged and ethically responsible partnerships with Indigenous communities.

## Methods

### Ethical approval and compliance

The study protocol for sample collection of Yanomami people was approved by the Institutional Research Ethics Committee (CEP), the National Research Ethics Commission (CONEP; no. 08069019.4.0000.5594) and by the Yanomami population of interest during local Kurikama assembly, for the period in which the samples were collected (September 2023). The Belgian sample collection was approved by the Commission for Medical Ethics U(Z) Ghent (CME; study reference ONZ-2023-0659). Human participants were selected based on health status, with the lack of any clinical signs of chronic or acute disease or discomfort, and no recent history of antibiotics use (up to 3 months prior to sampling). Informed consent was obtained from all participants, following procedures appropriate to their cultural context.

Animal experiments were approved by the University of Ghent Animal Ethics Committee and the Ethical Committee for Animal Experimentation at Ghent University’s Faculty of Medicine and Health Science (ECD 21-30 aanp).

### Metagenomic sequencing human samples

#### Human sample collection

Human stool samples were collected fresh using sterile sampling bags (VWR®, VWRI129-9024, Yanomami) or Fecotainer® (460869, Belgian) and aliquoted immediately into sterile 50ml falcons. Yanomami stool samples were kept at 4°C during the sample expedition and maintained in an anaerobic environment using the BD GasPak™ EZ Anaerobe Pouch System (#11777194) until shipment at -80°C to Belgium. Belgian stool samples were self-collected and stored at 4°C before transfer to the lab and storage at -80°C.

#### DNA Extraction and Sequencing

DNA extraction and shotgun metagenomic sequencing were performed by cmbio (USA). Genomic DNA was extracted from faecal material using the DNeasy PowerSoil Pro Kit (QIAGEN) according to the manufacturer’s instructions. DNA concentration was quantified fluorometrically using the Qubit system (Thermo Fisher Scientific). Libraries were sequenced as paired-end 2×150 bp reads on the Element AVITI and Illumina NovaSeq X Plus platforms.

#### Read-based taxonomic classification of human stool samples

Data was processed using nf-core/taxprofiler v1.2.3^55^ of the nf-core collection of workflows^56^, utilising reproducible software environments from the Bioconda^57^ and Biocontainers^58^ projects.

The pipeline was executed with Nextflow v24.10.2^59^. Tools used in the workflow included: Sequencing quality control with FastQC^60^. Short read preprocessing was performed using fastp^61^. Low-complexity sequence filtering was carried out using BBDuk^62^. Human host reads were removed by alignment to the GRCh38 reference genome using Bowtie2^63^ and SAMtools^64^. Taxonomic classification was performed using Kraken2^65^ against the PlusPF database (k2_pluspf_20241228^66^) obtained from the Kraken2 AWS index repository, using a confidence threshold of 0.1. Species abundances were estimated using Bracken^67^ with a read length parameter of 150 bp. Visualisation of results, where supported, was performed with Krona^68^. Pipeline results statistics were summarised with MultiQC^69^.

Downstream analyses were performed in R (v4.6.0) using Bracken-derived abundance estimates. Microbiome data handling and analysis were conducted using the mia package (v1.20.0) within the miaverse framework, following Orchestrating Microbiome Analysis with Bioconductor^70^. For differential abundance analysis, taxa were aggregated at the family level and analysed using MaAsLin2 (v1.26.0)^71^, applying a prevalence filter of 0.1 and a false discovery rate threshold (q ≤ 0.05). Data visualisation was performed using ggplot2 (v4.0.3)^72^.

#### Data availability

Individual-level metagenomic data generated in this study are available upon request, respecting Indigenous data sovereignty and ownership and complying with applicable Brazilian regulations governing genetic resources and benefit sharing. Requests should outline the proposed uses and scientific objectives, explain how data security and participant privacy will be protected, and describe the potential benefits for the Yanomami community.

### Human faecal microbiota transplant to GF mice

#### Donor selection for transplantation

To ensure that we did not select the most divergent samples of each cohort, and instead picked samples that were more centrally positioned within the Belgian and Yanomami cohort, Bracken-derived species abundance estimates were used. For each group, a reference composition was constructed by aggregating species-level counts across all samples within the group and normalizing the resulting profile to relative abundances. Individual samples were then compared to their group-level reference using Bray–Curtis dissimilarity to quantify compositional divergence, with lower values indicating greater similarity to the group profile. Samples above the 75^th^ percentile of within-cohort dissimilarity were excluded from donor selection for *in vivo* experiments. From the retained samples, donors were selected based on sample availability and covariate-matching constraints. Selected donor sets and associated experiments are listed in Table 1 and highlighted in red on Fig. 1d.

#### Preparation of faecal gavage mixture

Human stool sample (1g) was resuspended in 15ml pre-reduced PBS in an anaerobic environment (Concept 400 anaerobic workstation). Following resuspension, the sample was left for insoluble sediment to settle. The remaining supernatant was aliquoted in sterile CRYO.S tubes (Greiner, 123263), snap frozen and stored at -80 °C.

#### High-fat diet (HFD)-induced metabolic disease model

GF C57BL/6 (in-house derived colony) male mice were obtained from the Germ-Free and Gnotobiotic Mouse Facility (GFMF) Ghent, Belgium (LA2400637/LA1400637; ECD 24-72K). The mice were weaned at 3-4 weeks old and transferred to positive-pressure-Isocages (Isocage-P-Tecniplast) prior to colonization, under a 12h light/dark cycle, with controlled temperature (20-24°C) and relative humidity (45-65%). All experimental group allocations were random. Mice with the same donor microbiota were housed in the same cage, except for experiments with larger sample sizes using donors BEL_051, BEL_026 and YAN_009 where mice with the same donor microbiota were split into 2 cages, and YAN_013, where mice were put in 3 cages. No blinding of the investigators to the allocation was done during the experiments, since all experiments required gavages of known faecal donor material. All mice received 3 doses of faecal gavage mixture every 48h, maximum volume of 200μl/25g of body weight (20 gauge plastic feeding tube), to generate HMA mice. After a 2 week-period of colonization on standard chow (2018S, Teklad Global 18% Protein Rodent Diet, Envigo), the mice were given a high-fat diet (HFD, Bio-services, 60% fat diet D12492, E15742-34(7), irradiated) for 12 weeks. Mice had *ad libitum* access to food and water. Weekly weight and bi-weekly blood glucose measurements (Accu-Check® Guide meter) and stool collections were carried out. All mice handlings occurred in a sterile laminar flow to prevent contamination. A total of 7 human donor sets (Belgian vs. Yanomami) were used to colonize GF mice during HFD experiments (Table 1).

### HMA mice metabolic and immune response evaluation

#### Intraperitoneal glucose tolerance test (IPGTT)

An IPGTT test was conducted either on the day of sacrifice (BEL_051 vs. YAN_009 and BEL_026 vs. YAN_013), or 2 days prior to sacrifice (all other donors tested). Mice were fasted overnight (14 - 16h) before injecting a 20% (2g kg^-1^ body weight – BEL_051 vs. YAN_009) or 10% (1g kg^-1^ body weight - all other donors tested) glucose solution intraperitoneally. Blood was collected by tail prick at 0, 15, 30, 60 and 120 min after injection and glucose levels were measured using the Accu-Check® Guide meter.

#### Quantification of metabolic hormones and serum cytokines

Metabolic hormone concentrations in fasting serum were determined by Bio-Plex Pro Mouse Diabetes 8-Plex Assay (#171F7001M, Bio-Rad), according to manufacturer’s instructions.

Cytokine (IL-6, TNF, IL-17, IFNγ, MCP-1) levels were measured in the serum (non-fasting) of mice using the Bioplex® Multiplex Immunoassay System (Biorad), according to manufacturer’s instructions. Read-out was performed using the Bio-Plex 200 system (Bio-Rad). For experiments testing BEL_051 vs. YAN_009 and BEL_026 vs. YAN_013, serum was collected 120 minutes after glucose injection for IPGTT (1g kg^-1^ body weight, i.p., preceded by overnight fast), since fasting serum was all used for the quantification of metabolic hormones. For all other donor sets, non-fasting serum was collected.

#### Free fatty acid quantification

Free fatty acids were quantified in serum using the Sigma-Aldrich Free Fatty Acid Assay Kit (MAK466), according to manufacturer’s instructions. Serum samples were used undiluted and background corrections were made using a blank sample and a serum blank (serum with assay buffer, no working reagents added). For experiments testing BEL_051 vs. YAN_009 and BEL_026 vs. YAN_013, serum was collected 120 minutes after glucose injection for IPGTT (1g kg^-1^ body weight, i.p., preceded by overnight fast), since fasting serum was all used for the quantification of metabolic hormones. For all other donor sets, non-fasting serum was collected.

#### ALT activity quantification

Alanine aminotransferase (ALT) activity was determined in the serum of mice following 12 weeks of HFD using the Sigma-Aldrich ALT Activity Assay Kit (MAK571), according to manufacturer’s instructions. Serum samples were diluted 100x in ALT Assay Buffer and background corrections were made using a blank sample. For experiments testing BEL_051 vs. YAN_009 and BEL_026 vs. YAN_013, serum was collected 120 minutes after glucose injection for IPGTT (1g kg^-1^ body weight, i.p., preceded by overnight fast), since fasting serum was all used for the quantification of metabolic hormones. For all other donor sets, non-fasting serum was collected.

#### Histology

Liver, eWAT, colon and ileum samples were fixed in 4% PFA, embedded in paraffin and sectioned at 5 μm. All sections were stained with H&E and imaged on the Zeiss AxioScan Z.1 Digital Slide Scanner. Histopathological scoring of the liver was performed on H&E stained liver sections using a previously published protocol ^73^. One section per mouse was blindly scored for hepatocellular hypertrophy, micro- and macrovesicular steatosis, and number of inflammatory foci. Quantification of adipocyte size was performed on H&E-stained eWAT sections, using the Adiposoft plugin for ImageJ.

To quantify lipid associated macrophages (LAMs) in eWAT tissue, sections (5µm thick, FFPE) were dewaxed, followed by antigen retrieval in Antigen Unmasking Solution, Citric Acid Based (Vector Laboratories Inc., H-3300) using a PickCell 2100-Retriever. Following photobleaching (Merscope photobleacher, Vizgen, 1h, in 3% H_2_O_2_ + 20 mM NaOH (in PBS)), slides were blocked (1% donkey serum, 0.5% fish skin gelatine and 2% BSA in PBS, 1h, RT) and incubated with F4/80 (Invitrogen, 14-4801-81, 1/200) and GPNMB (Abcam, ab188222, 1/100) antibody (overnight, 4°C). Finally, slides were incubated with secondary antibodies donkey-anti-rabbit Alexa Fluor 568 (1/500) and donkey-anti-rat Dylight 755 (1/1000) to visualize GPNMB and F4/80, respectively, in combination with DAPI (1/1000, 2h, RT). Slides were mounted with Polyvinyl alcohol mounting medium with DABCO® (Merck, 10981) and sections were imaged on the Zeiss AxioScan 7 Digital Slide Scanner. Quantification of LAM density (cells/mm²) was performed on 3 regions of an eWAT section per mouse using QuPath-0.5.1.

Liver samples were snap frozen, embedded in Tissue-Tek® O.C.T. Compound (4583) and sectioned at 12 µm. For Oil Red O staining, cryosections were left to dry at RT for 15min before rinsing with PBS and fixing in 4% PFA at RT for 30 min. Sections were then washed with PBS, followed by a wash with 1,2-propanediol (Sigma-Aldrich 398039) for 5 min. Sections were then stained with a pre-warmed stock solution of Oil Red O (sigma O-0625, 0.5% stock solution in 1,2-propanediol) at 60°C. Hereafter, slides were destained in 85% 1,2-propanediol for 5 min at RT whilst shaking, followed by washing for 5min with running water and counterstaining with Mayer’s haematoxylin for 30sec at RT. After a final 5min wash with running water, slides were mounted with Polyvinyl alcohol mounting medium with DABCO® (Merck, 10981). Sections were imaged on the Zeiss AxioScan Z.1 Digital Slide Scanner and measurement of average lipid droplet size was performed on 3 Oil Red O stained sections per mouse using QuPath-0.5.1.

Colon samples containing faecal matter were fixed in Carnoy fixative (60% methanol, 30% chloroform and 10% glacial acetic acid), embedded in paraffin and sectioned at 5 µm. The sections were stained with AB-PAS by incubating dewaxed slides in 1% Alcian Blue for 20 min. Sections were then washed with water and incubated with 1% periodic acid (Sigma-Aldrich, 395132) for 15 min followed by Schiff’s reagent (Sigma-Aldrich, 395-2-016) for 10 min. Finally, sections were counterstained with Mayer’s haematoxylin for 30 s before washing, dehydration and mounting with Entellan (Merck). Following imaging on the Zeiss AxioScan Z.1 Digital Slide Scanner, mucus thickness was measured using the Zeiss Zen 3.9 software. Average mucus thickness was calculated per sample based on 5 measurements per section.

#### Cell isolation and flow cytometry analysis of liver and eWAT

Prior to liver dissection, liver perfusion was performed with 5ml PBS, after which the liver was excised and kept in RPMI Medium 1640 (gibco, 52400-025) on ice. Liver cell isolation was performed according to published protocol^74^.

Epididymal fat pad (eWAT) cell isolation was performed by digesting eWAT tissue in RPMI + Collagenase II (1mg/ml, Worthington, LS004176) for 20-30 minutes (37°C, 150 rpm). Following digestion, EDTA was added to a final concentration of 10mM (UltraPure^TM^ 0.5M EDTA, Invitrogen, 15575-038) for 3 minutes. Digestion was neutralized by adding RPMI + 10% FBS (heat inactivated, LPS-, Serena) in equal volume. After filtering through a 100µm strainer, cells were centrifuged (500 x g, 7 min, RT) and the cell pellet was resuspended in ACK lysing buffer (gibco, A10492-01, 5 min, 4°C). ACK lysis was neutralized by adding RPMI + 10% FBS (10ml), cells were filtered using a 70µm filter, centrifuged (500 x g, 7 min, 4°C), and the final cell pellet was resuspended in PBS + 2% FBS + 2mM EDTA.

For both tissues, an innate immune (myeloid) specific and T cell specific panel were analysed.

In the myeloid panel, following cell subsets were identified: neutrophils (CD45^+^CD3^-^CD19^-^NK1.1^-^Ly6G^+^), monocytes (CD45^+^CD3^-^CD19^-^NK1.1^-^Ly6G^-^Ly6C^+^), DCs (CD45^+^CD3^-^CD19^-^NK1.1^-^Ly6G^-^Ly6C^-^F4/80^-^CD64^-^CD11c^+^MHCII^+^), macrophages (CD45^+^CD3^-^CD19^-^NK1.1^-^Ly6G^-^Ly6C^-^F4/80^+^CD64^+^), infiltrating macrophages (CD45^+^CD3^-^CD19^-^NK1.1^-^Ly6G^-^Ly6C^-^F4/80^+^CD64^+^VSIG4^-^), resident macrophages (CD45^+^CD3^-^CD19^-^NK1.1^-^Ly6G^-^Ly6C^-^F4/80^+^CD64^+^VSIG4^+^), pro-inflammatory or alternatively activated immunoregulatory infiltrating and resident macrophages were defined as iNOS^+^ or ARG1^+^, respectively.

In the T cell panel, following cell subsets were identified: T cells (CD45^+^CD3^+^), cytotoxic T cells (CD45^+^CD3^+^CD8^+^CD4^-^), T helper cells (CD45^+^CD3^+^CD8^-^CD4^+^), regulatory T (Treg) cells (CD45^+^CD3^+^CD8^-^CD4^+^Foxp3^+^), Th1 cells (CD45^+^CD3^+^CD8^-^CD4^+^Tbet^+^), Th2 cells (CD45^+^CD3^+^CD8^-^ CD4^+^GATA3^+^) and Th17 cells (CD45^+^CD3^+^CD8^-^CD4^+^RORγt^+^).

All antibodies used for flow cytometry analysis and used gating strategy are described in Suppl. Table 1 and Suppl. Fig. 4, respectively. All samples were fixed and permeabilized using eBioscience™ Foxp3 / Transcription Factor Staining Buffer Set (Invitrogen, cat. no. 00-5523-00) prior to intracellular staining with iNOS/ARG1 for the myeloid panel or Foxp3/Tbet/GATA3/RORγt for the T cell panel. Cell acquisition was done on the BD FACSDiscover™ A8 Cell Analyzer, whereby the autofluorescence of the specific tissue (liver and eWAT) was taken into account using an unstained control. Settings on the BD FACSDiscover™ A8 Cell Analyzer were kept identical across experiments and relative frequencies are reported to ensure cross-experiment comparability.

A conjugate hierarchical generalized linear model (HGLM) was used to analyse immune cell proportion data from each flow cytometry panel. The model was fitted using the hglm package (v2.2-1 ^75^) in R (v4.3.1 ^76^). For the pooled analysis, the model included Experiment and Group as fixed terms, with Donor as a random term. For per-experiment analyses, only Group was included as a fixed term, with Donor as a random term. Fixed effects followed a Binomial distribution, while random effects used a Beta distribution, both utilizing a logit link function. The variance dispersion parameter for responses was estimated via extended quasi-likelihood (EQL). T-statistics assessed the significance of group effects on the logit-transformed scale, with *P* values derived from a t-distribution. Fixed effect estimates were exponentiated to obtain odds ratios with 95% confidence intervals. Group effects are illustrated in forest plots showing results on both the logit-transformed scale and the odds ratio scale, generated using ggplot2 (v4.0.3 ^72^).

#### Gene expression analysis

Colon and eWAT tissues were harvested from mice following 12 weeks of HFD and stored in RNAprotect Tissue Reagent (Qiagen, Cat no. 76104) at -20°C. Tissue was homogenized in TRIzol® (1ml, Invitrogen, 15596018), using the TissueLyser II (Qiagen). Chloroform was added (200µl) and samples were centrifuged (12 000 x g, 15 min, 4°C) after vigorous shaking and incubation (3 min, RT). The upper aqueous phase was mixed with isopropanol (500 µl), incubated (10 min, RT) and centrifuged (12 000 x g, 10 min, 4°C). The pellet was then washed in ethanol (75%, 1 ml), followed by centrifugation (7500 x g, 5 min, 4°C), air drying of the pellet and finally resuspension in DEPC-treated and nuclease-free water (fisher bioreagents, BP561-1). RNA concentration and quality was measured using the NanoPhotometer® N120 (Implen). cDNA synthesis was performed using the QuantiTect® Reverse Transcription Kit (Qiagen, cat no. 205313), according to manufacturer’s instructions. RT-qPCR reactions were performed with 10 ng of cDNA in a total volume of 10µl with SensiFAST^TM^ SYBR® No-ROX (Bioline) on the LightCycler 480 (Roche), in triplicate. A list of used primers can be found in Suppl. Table 2. Data was analysed using qbase+ software.

### Liver transcriptome analysis

#### Sample preparation

Liver tissue was harvested from mice following 12 weeks of HFD (n = 8/group) and stored in RNAprotect Tissue Reagent (Qiagen, Cat no. 76104) at -20°C. Tissue was homogenized in TRIzol® (1ml, Invitrogen, 15596018), using the TissueLyser II (Qiagen). Chloroform was added (200µl) and samples were centrifuged (12 000 x g, 15 min, 4°C) after vigorous shaking and incubation (3 min, RT). The upper aqueous phase was mixed with isopropanol (500 µl), incubated (10 min, RT) and centrifuged (12 000 x g, 10 min, 4°C). The pellet was then washed in ethanol (75%, 1 ml), followed by centrifugation (7500 x g, 5 min, 4°C), air drying of the pellet and finally resuspension in DEPC-treated and nuclease-free water (fisher bioreagents, BP561-1). RNA concentration and quality was measured using the NanoPhotometer® N120 (Implen).

#### Library preparation and sequencing

Quality control, library preparation and subsequent sequencing was performed by Novogene. For library preparation, the Novogene NGS Stranded RNA Library Prep Set (PT044) was used. Sequencing was performed on the NovaSeq X Plus Series platform.

#### RNA-seq processing and quality control

Raw sequencing data were processed using nf-core/rnaseq v3.24.0^77^ from the nf-core collection of workflows^56^, utilising reproducible software environments provided by Bioconda^57^ and Biocontainers^58^. The pipeline was executed using Nextflow v25.04.6^59^. Transcripts were mapped to the mouse reference genome GRCm39 (GCA_000001635.9) using Ensembl annotation release 112. Sequencing and alignment quality metrics were assessed using the quality-control tools implemented within the nf-core/rnaseq workflow using default settings.

#### RNA-seq downstream analysis

Salmon transcript abundance estimates generated by the pipeline were imported into R (v4.6.0) using tximport (v1.40.0)^78^ and summarised to gene-level counts for downstream analyses. The experimental design and sample relationships were assessed using principal component analysis (PCA) performed on variance-stabilized count data generated using DESeq2 (v1.52.0)^79^. Differential gene expression analysis was conducted using DESeq2. Genes with an adjusted *P* value ≤ 0.05 and an absolute log2 fold change ≥ 0.5 were considered differentially expressed.

Gene set enrichment analysis (GSEA) and over-representation analysis (ORA) were performed using the clusterProfiler package (v4.20.0)^80,81^. Gene set annotations were retrieved from the msigdbr package (v26.1.0)^82^. The following MSigDB^83–86^ collections were included: Gene Ontology terms: Biological Processes and Molecular Functions (C5); canonical pathways from KEGG (CP:KEGG_LEGACY and CP:KEGG_MEDICUS), Reactome (CP:REACTOME), WikiPathways (CP:WIKIPATHWAYS), and chemical/genetic perturbations (CGP) (C2); Hallmark gene sets (H); and transcription factor target gene sets from the GTRD subcollection (C3).

For GSEA, differential gene expression (DGE) results were shrunk using the ashr package (v2.2.63)^87^ to reduce variance associated with lowly expressed genes. A preranked gene list was constructed by ordering genes based on their log2 fold change. Only gene sets containing between 15 and 500 genes^88^ were included to ensure statistical robustness and biological interpretability. Significantly enriched pathways were defined by *P* value < 0.05 and *q-*value < 0.2 for ORA, and *P* value < 0.05 for GSEA.

Transcription factor activity was inferred using the decoupleR package (v2.17.0)^89^. Figures were generated using ggplot2 (v4.0.3).

#### Data availability

Bulk RNA-seq data (raw FASTQ files and processed count matrices) have been submitted to the NCBI Gene Expression Omnibus (GEO) under accession number GSE350965.

### Metagenomic analysis mouse gut microbiota

#### Sample collection, DNA extraction, and sequencing

Faecal samples were collected from recipient mice before and after HFD administration. In the experiments comparing BEL_051 vs. YAN_009 and BEL_026 vs. YAN_013, individual mouse samples were sequenced, whereas other experiments were represented by faecal pools generated for each human donor and time point. DNA was extracted from faecal mass by Novogene using Tiangen Magnetic Universal Genomic DNA Kit, following the procedures recommended by the manufacturer. Extracted DNA was assessed using the Agilent Fragment Analyzer 5400 system with the Agilent Genomic DNA Analysis Kit (DNF-488) to evaluate DNA integrity and size distribution. DNA concentration was measured using a Qubit Fluorometer with the Qubit dsDNA HS Assay Kit. Shotgun metagenomic libraries were prepared using the Novogene NGS DNA Library Prep Set (Cat No.PT004) and sequenced on the Illumina NovaSeq X Plus platform using paired-end, 2×150bp sequencing layout. DNA extraction, library preparation, and sequencing procedures were conducted by Novogene (Beijing, China). Gavage inoculum samples were processed using a similar procedure. Details regarding the sequencing and processing of donor metagenomics data can be found in the section above (Metagenomic analysis human samples).

#### Read preprocessing and metagenomic assembly

All bioinformatic procedures used default parameters unless otherwise stated. Initial processing was performed using modules 1 and 3 of the nf-core/mag pipeline v5.4.2^90^. Sequencing adapters, poly-G tails, and low-quality reads were removed with fastp^61^. Reads aligning to mouse (GRCm39) and PhiX reference sequences were removed with Bowtie2^63^ using “--host_removal_verysensitive” parameter. Filtered reads were co-assembled separately for each human microbiota donor using the nf-core/mag “--coassemble_group” option. Assemblies were generated with MEGAHIT^91^. Sequencing, assembly, read-mapping, gene-prediction, and annotation quality metrics for each donor-specific co-assembly are summarized in Suppl. Table 3.

#### Taxonomic assignment, functional annotation and feature quantification

Assembled contigs were taxonomically classified using MMseqs2 taxonomy v16.747c6^92^ against the Genome Taxonomy Database (GTDB) v226. Contigs not assigned to Bacteria or Archaea were subsequently queried against the NCBI non-redundant protein database (downloaded on 18 July 2025). Full taxonomic lineages were generated using --tax-lineage 1 and reformatted with TaxonKit v0.20.0^93^ using reformat2. Protein-coding sequences were predicted from each donor-level co-assembly using Prokka v1.14.6^94^ with the “--metagenome” option. Predicted proteins were functionally annotated with eggNOG-mapper v2.1.12^95^ against eggNOG v5.0.2 using DIAMOND in ultra-sensitive mode. KEGG Orthology (KO) identifiers, their corresponding KEGG pathway, and higher-level annotations were retained for downstream analyses. Filtered reads were mapped to assembled contigs using Kallisto v0.46.1 ^96^ in quant mode. Contig abundance was estimated by length-normalized counts per million (CPM). Predicted genes inherited the abundance of their parent contig.

Count matrices and annotations were imported into R v4.3.2^76^. Counts were collapsed to non-redundant taxonomic/KO identifiers for taxonomic and functional analyses, respectively. Features were retained when detected in at least 10% of samples within each donor group. The main results used individual recipient-mouse profiles from the BEL_026 versus YAN_013 donors. For analysis across all experiments, one profile was generated per human donor and time point. For experiments in which multiple mouse samples were sequenced, feature abundances were averaged across recipient mice.

For other experiments, the profile obtained from the pooled faecal sample was used directly. This produced seven Belgian and seven Yanomami donor-level profiles at both the pre- and post-HFD time points.

#### Taxonomic profiling, microbiota engraftment and ecological analysis

Taxonomic analyses were conducted using phyloseq v1.46.0^97^ and utility functions from metagMisc v0.5.0^98^ and microbiome v1.24.0^99^ R packages. For taxonomic profile and alpha diversity analysis, genera with a relative abundance below 0.1% within a sample were treated as absent, after which genera present in more than 10% of samples were retained. Furthermore, features not classified down to the genus level were disregarded. For feature-level beta-diversity analyses, a minimum relative abundance threshold of 0.01% was applied before the same prevalence filter. FMT engraftment was calculated at the genus level. For each human donor, genera were ranked by relative abundance, and the genera cumulatively accounting for 99% of the donor microbiota were defined as the donor core. Engraftment was calculated as the proportion of donor-core genera detected in the corresponding recipient-mouse donor-level profile.

Shannon and Gini–Simpson indices were calculated from genus-level profiles using the alpha function (microbiome package). Bray–Curtis dissimilarities were calculated from filtered unaggregated taxonomic features in phyloseq and visualized by principal coordinate analysis (PCoA). Differences in community composition were evaluated by PERMANOVA using adonis2 in vegan v2.7-2^100^. Tests used 999 permutations restricted by cage as blocking strata (only main comparison). Pairwise comparisons among combined donor-cohort and time-point groups were performed using pairwise.adonis2 from pairwiseAdonis v0.4.1^101^ with FDR correction. Homogeneity of multivariate dispersion was assessed using betadisper, and distances to group centroids were compared. Pre- to post-HFD compositional or functional shifts were calculated as the Bray–Curtis dissimilarity between samples obtained from the same mouse/donor group at the two time points. Distances were compared between Belgian- and Yanomami-derived communities.

#### Functional profiling

Functional profiles were analysed at the KO feature and KEGG metabolism pathway levels. Because individual KOs may belong to multiple pathways or subclasses, their abundance was distributed across all applicable categories and then TSS-normalized. Bray–Curtis dissimilarities and other subsequent analyses involving this metric were calculated as previously described. For pathway contribution plots, the abundance of KOs assigned to each pathway was partitioned according to the genus assigned to the corresponding contig. Each genus contribution was calculated as its proportion of the total pathway-associated KO abundance.

#### Differential-abundance and statistical analysis

Differential abundance of genera and KOs was assessed using MaAsLin2 v1.16.0^71^. Time point or donor cohort was included as the fixed effect, depending on the contrast. Associations were considered significant at an absolute regression (β) coefficient greater than 1 and a BH-adjusted *q-*value below 0.05. Associations between genus abundances and host metabolic, inflammatory, and histopathological measurements were evaluated using separate MaAsLin2 models for each host variable, with each host measurement included as a fixed effect. Associations with BH-adjusted *q* < 0.1 were retained for visualization. For the analysis across all donors, host measurements were summarized at the donor level using mean values. Pathway-level differences were identified using LEfSe, implemented with run_lefse in microbiomeMarker v1.13.2^102^. Pathways with an absolute LDA score ≥2 and an FDR-adjusted *P* value <0.05 were considered significantly associated. To assess the robustness of these results using a compositional differential-abundance method, pathway-level associations were additionally evaluated using ANCOM-BC, implemented with run_ancombc in microbiomeMarker (FDR-adjusted *P* value <0.05). Before visualization, significantly associated KEGG pathways were manually curated to exclude pathway labels that were not directly or indirectly attributable to bacterial metabolism, to avoid interpreting shared or overlapping bacterial KOs as evidence for predominantly host-, plant-, or other non-bacterial biological processes.

General differences among groups were assessed using the Kruskal–Wallis test, followed, when significant, by pairwise Wilcoxon rank-sum tests with Benjamini–Hochberg correction to control the False Discovery Rate (FDR). FDR-adjusted *P* values < 0.05 were considered statistically significant, whereas values between 0.05 and 0.1 were interpreted as trends. For heatmap visualization, relative abundances were centred log-ratio transformed and subsequently Z-scaled separately for each feature across profiles. The 50 genera or metabolic pathways with the greatest absolute MaAsLin2 coefficients across the tested contrasts were selected. Rows and columns were hierarchically clustered using Euclidean distance and complete linkage.

#### Data availability

Metagenomic shotgun sequencing data from mouse stool samples have been submitted to the NCBI Sequence Read Archive (SRA) under a BioProject, with BioSample metadata. The submission is in progress, and the accession numbers (PRJNA, SRR) will be provided upon assignment and added to the manuscript.

### Statistical analysis

Data from the highlighted experiment comparing BEL_026 vs. YAN_013 are shown in box plots, whereby the minimum to maximum is shown with all replicates included, the horizontal line within the box shows the median, and the box limits extend from the 25th to 75th percentile. Exact *P* values, effect sizes and confidence intervals are mentioned in the accompanying figure legend. Normality and homoscedasticity of residuals was evaluated using standard diagnostic plots. Data was analysed using a Welch’s t test in case of normally distributed data. Data that did not fit the assumption of normality was analysed using a Mann-Whitney test. These analyses were performed using GraphPad Prism 11. Weight follow up was analysed using a linear mixed-effects model fitted by restricted maximum likelihood (REML), with a quadratic effect of time (weight ∼ GROUP * (TIME + I(TIME^2)) + (TIME | ID)). Group, time, and their interaction are included as a fixed effect, with mouse ID as a random effect. Time was modelled as a quadratic term, since this provided a substantially better fit than a linear model. No correction for start weight was performed, since body weight measurements started at week 0, 2 weeks after microbiota colonization, which is considered a post-intervention measurement rather than a true baseline. We recognize that this is a limitation of the study and therefore include body weight measurement before microbiota colonization in the following experiments.

Data from pooled experiments was analysed using a linear mixed-effects model fitted by REML (Outcome_Metric ∼ Treatment + (1 | Donor)). Microbiota origin (i.e. Belgian vs. Yanomami, Treatment effect in model) was included as a fixed effect, and donor was modelled as a random intercept to account for non-independence of mice derived from the same donor. Because each donor contributed animals to only one treatment group, donor was nested within treatment by design. The model partitions variability into between-donor and within-donor components, thereby accounting for clustering while estimating treatment effects at the cohort level. Model assumptions (normality and homoscedasticity of residuals) were evaluated using standard diagnostic plots. Weight follow-up from the pooled experiments was analysed using linear mixed-effects model fitted by REML with fixed effects for Experiment, Time, Group, and Time x Group interaction, corrected for start weight, and random effects for mouse ID (repeated measures), donor, and donor–time interaction.

All statistical analyses performed during flow cytometry analysis, bulk mRNA sequencing analysis and metagenomic analysis are mentioned in the accompanying method sections.

### Use of AI

Generative artificial intelligence (Claude (Anthropic) and Microsoft Copilot) was used to support the research process by assisting with literature exploration, text editing, language refinement, summarization of source materials provided by the authors, and code drafting during manuscript preparation. All scientific judgments, data interpretation, critical evaluation of sources, and final manuscript content were performed, verified, and approved by the authors. The AI system was not involved in data collection and is not considered an author.

## Acknowledgements

We thank all study volunteers for their contribution to this research; the Germ-Free and Gnotobiotic Mouse Facility Ghent (GFMF) and Vanessa Andries for coordinating the germ-free and gnotobiotic mouse experiments, and Korneel Barbry for animal care; Katrien Van der Borght, Julie Van Duyse, Elien Ruyssinck and Sofie Van Gassen of the VIB Flow Core Ghent, Amanda Goncalves of the VIB BioImaging Core Ghent and Kelly Lemaire for experimental support; Els Adriaens and Marnik Vuylsteke for statistical consultancy; Dirk Van Dijck for logistical support during the sampling expedition in Brazil; and COPAN for sponsoring sampling material. Marie Thorp is a predoctoral fellow supported by an FWO doctoral fellowship (11A1823N, 11A1825N). Research performed in the Vereecke lab was supported by FWO (3G0H2522, G0APB25N) and Ghent University (bof/baf/4y/2024/01/269, 01G03122, Yanobiome). Research performed in the Paulino lab was supported by The São Paulo Research Foundation (FAPESP) and FWO (2024/08541-5, 2025/08894-8, 2026/02527-6). Research performed in the Allen-Vercoe lab was supported by the Canada Research Chairs program (CRC-2025-00071).

## Author contributions

M.T. and L.V. conceived the study and wrote the manuscript with input from L.A.L.d.C. and L.C.P.. M.T. designed and performed all *in vivo* experiments, with assistance of G.B.. L.A.L.d.C. and M.C. performed bioinformatic analysis on the human sequencing data, and L.A.L.d.C. performed all bioinformatic analysis of the *in vivo* experiments. V.A. coordinated all germ-free and gnotobiotic mouse experiments. M.J. provided input and feedback on experimental setup. S.G.S. performed histopathological scoring of mouse liver sections. B.D.P. provided input and feedback on statistical analysis and experimental setup. M.T., C.C., D.G. and L.C.P. co-organized and performed the sampling expedition in Brazil, M.T. organized and performed sampling in Belgium. G.v.L., E.A.V., L.C.P. and L.V. provided ideas and feedback on the project. All authors reviewed and approved the final manuscript.

## Notes

### Competing Interest Statement

The authors have declared no competing interest.

