## Supplementary figures and tables for "Indigenous Yanomami gut microbiota protects against diet-induced metabolic dysfunction"

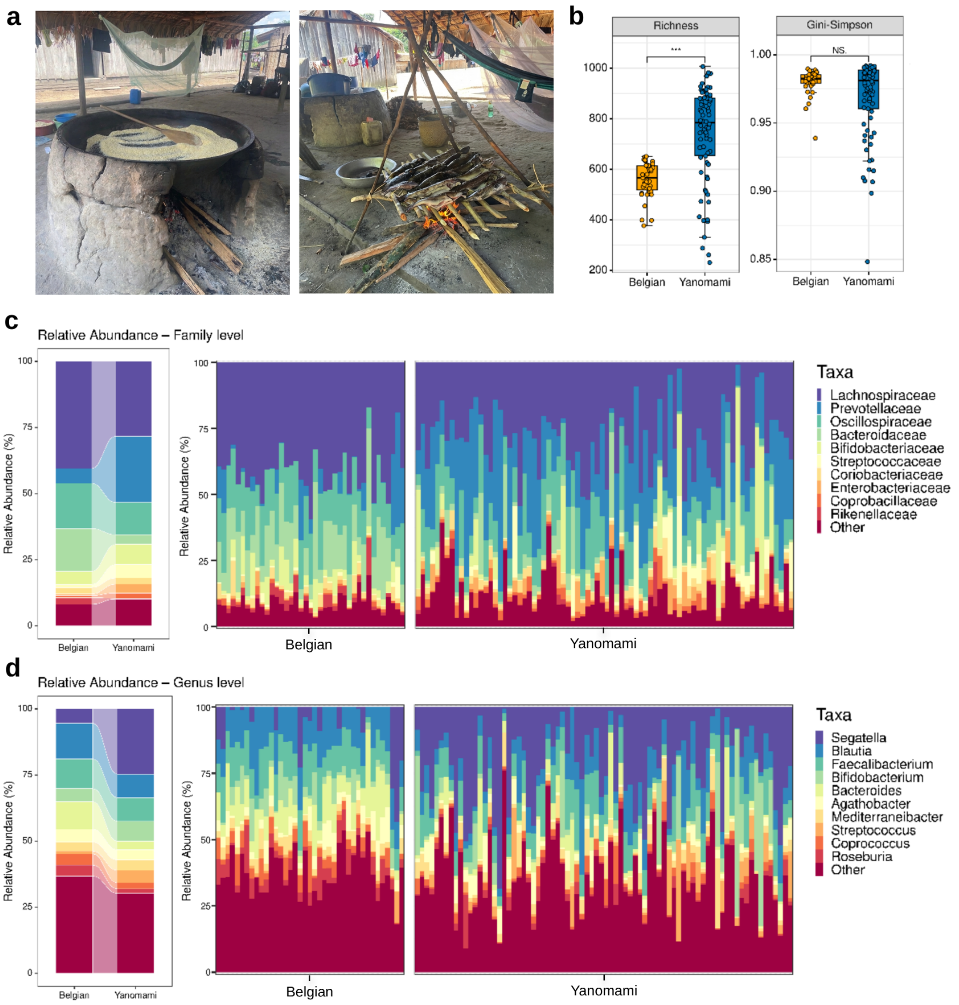


***Suppl. Fig. 1|Belgian and Yanomami cohorts differ in gut microbiota composition. a,*** *Examples of the high fibre, low fat diet of Yanomami people, with high consumption of ground cassava root (left), plantain, legumes, fish (right) and occasional meat.* ***b,*** *Distribution of gut microbiota alpha diversity in Belgian (n = 39) and Yanomami (n = 78) cohorts. Richness and Gini-Simpson values were calculated at the taxonomic feature level from contig-based classifications, using the best taxonomic assignment up to the species level. Statistical significance between groups was assessed using the Wilcoxon test (Richness: p-adj < 0.001; Gini-Simpson: p-adj = 0.761).* ***c,*** *Relative abundance at the family level, summarized for Belgian vs. Yanomami cohorts (left) or for individual donors (right).* ***d,*** *Relative abundance at the genus level, summarized for Belgian vs. Yanomami cohorts (left) or for individual donors (right).*

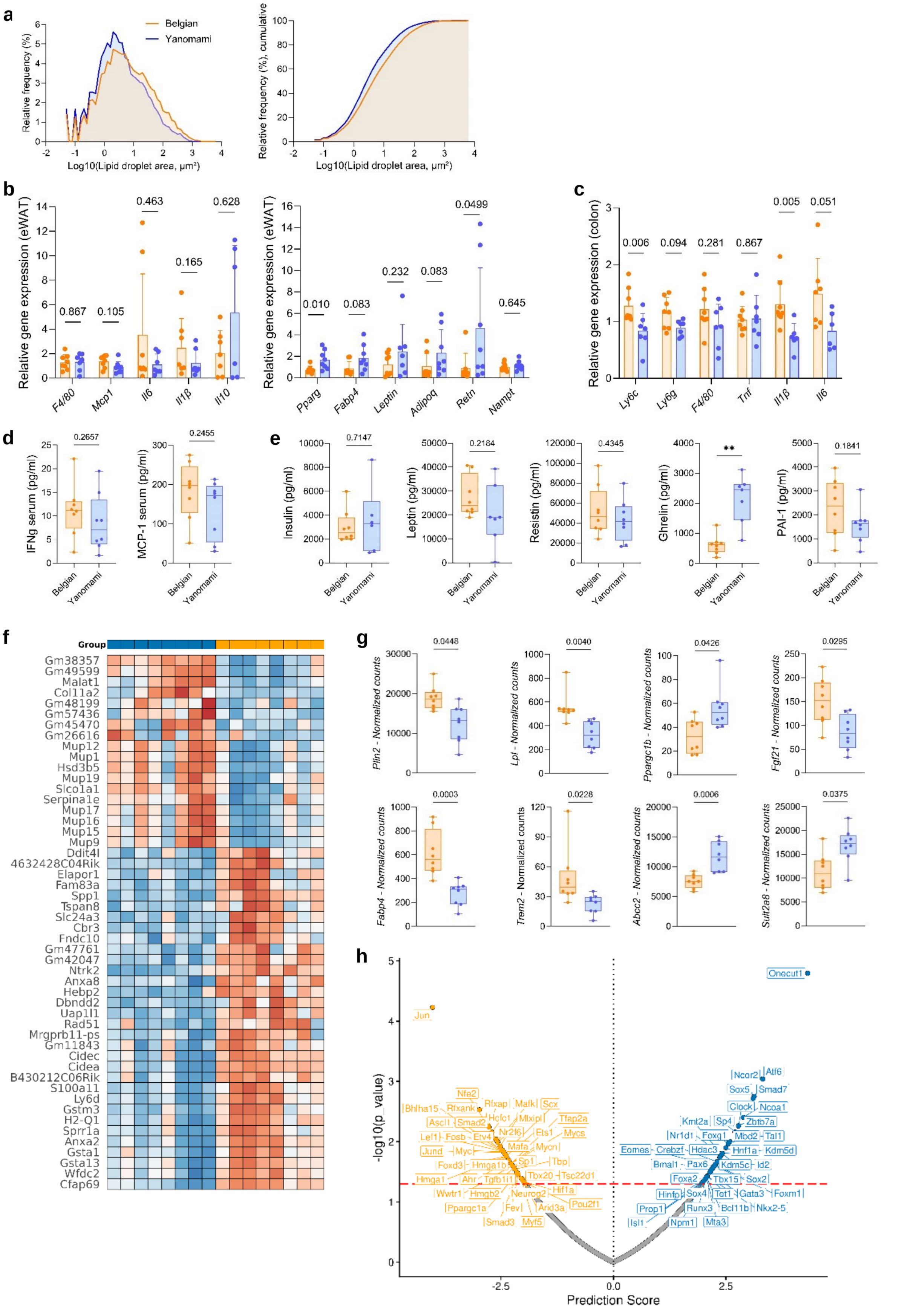


***Suppl. Fig. 2|Altered inflammatory and metabolic phenotype in mice colonized with a Belgian vs. Yanomami gut microbiota upon HFD. a,*** *Histogram showing the distribution of lipid droplets as the relative frequency (left) or cumulative frequency (right) of log10(lipid droplet area, µm²).* ***b,*** *Gene expression analysis of markers related to inflammation (left) or metabolism and adipokines (right) in eWAT tissue of mice following 12 weeks of HFD. Gene expression was quantified relative to Ywhaz and Rplp0 expression levels and scaled to an average value of 1. Significance was*

*determined using a Mann-Whitney test (exact two-tailed P values indicated).* ***c,*** *Gene expression analysis of markers related to inflammation in colon tissue of mice following 12 weeks of HFD. Gene expression was quantified relative to Gapdh and β-actin expression levels and scaled to an average value of 1. Significance was determined using a Mann-Whitney test (exact two-tailed P values indicated).* ***d****, Inflammatory cytokines measured in serum of mice following 12 weeks of HFD (non-fasting, 120min post glucose injection for IPGTT). Significance was determined using a Mann-Whitney test (IFNγ: median 11.20 (Belgian, n=8) vs. 7.065 (Yanomami, n=8), Hodges-Lehmann difference = −2.995, Mann-Whitney U = 21, exact two-tailed P = 0.2657; MCP-1: median 196.9 (Belgian, n=8) vs. 172.1 (Yanomami, n=8), Hodges-Lehmann difference = −32.27, Mann-Whitney U = 20.50, exact two-tailed P = 0.2455).* ***e,*** *Diabetes markers measured in the fasting serum of mice following 12 weeks of HFD (16h fasting period). Significance was determined using a Welch's t test (insulin: t(6,799) = 0.3811, P = 0.7147, mean difference 477.2, 95% CI −2501 to 3456; leptin: t(10,27) = 1.311, P = 0.2184, mean difference -7488, 95% CI −20 168 to 5193; resistin: t(13,61) = 0.8053, P = 0.4345, mean difference -9196, 95% CI −33 754 to 15 362; ghrelin: t(7,644) = 4.743, P = 0.0017, mean difference 1534, 95% CI 781.9 to 2284; PAI-1: t(11,75) = 1.411, P = 0.1841, mean difference -707.8, 95% CI -1803 to 387.6).* ***f-h,*** *Bulk mRNA sequencing of liver tissue from Belgian vs. Yanomami mice following 12 weeks of HFD.* ***f****, Heatmap showing the top 50 differentially expressed genes in the liver of Belgian- vs. Yanomami-HMA mice following 12 weeks of HFD.* ***g,*** *Boxplots showing normalized expression of specific DEGs in liver of Belgian- and Yanomami HMA-mice. DEGs were determined using DESeq2 and were considered differentially expressed in case of an adjusted P value ≤ 0.05 and an absolute log2 fold change ≥ 0.5.* ***h,*** *Volcano plot of transcription factor (TF) activity scores inferred using decoupleR. Differential TF activity between Yanomami- and Belgian-HMA liver samples was estimated using the ULM method implemented in decoupleR with the CollecTRI regulatory network. The x-axis shows the difference in inferred TF activity scores between groups, and the y-axis shows statistical significance (−log10 P value). The dashed horizontal line indicates the significance threshold (P < 0.05). All significantly differentially regulated TFs are labelled.*

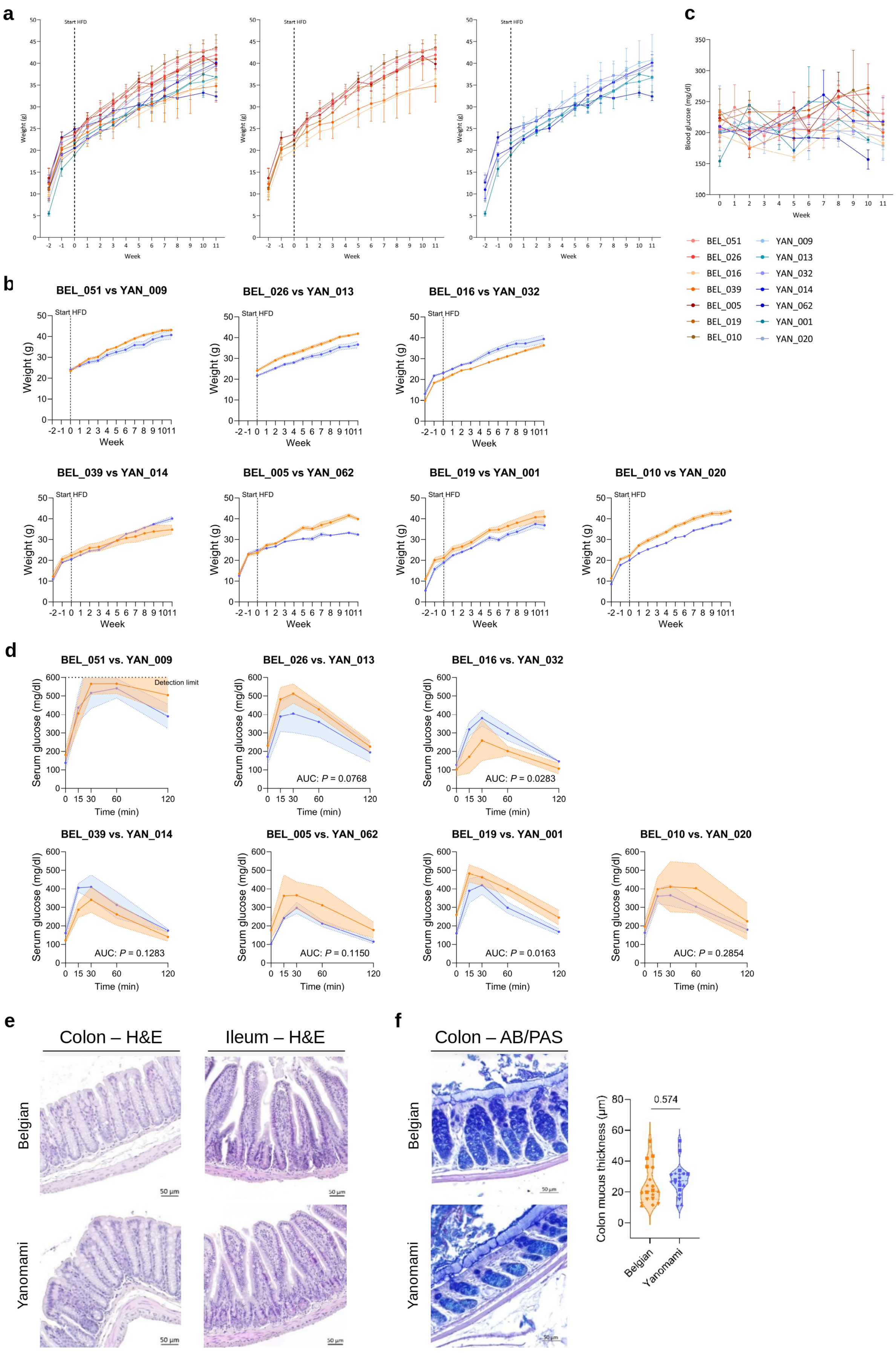
a
Start HFD
Weight (g)
Week
c
Blood glucose (mg/dl)
Week
BEL_051
BEL_026
BEL_016
BEL_039
BEL_005
BEL_019
BEL_010
YAN_009
YAN_013
YAN_032
YAN_014
YAN_062
YAN_001
YAN_020
b
BEL_051 vs YAN_009
BEL_026 vs YAN_013
BEL_016 vs YAN_032
BEL_039 vs YAN_014
BEL_005 vs YAN_062
BEL_019 vs YAN_001
BEL_010 vs YAN_020
d
BEL_051 vs. YAN_009
Detection limit
BEL_026 vs. YAN_013
AUC: P = 0.0768
BEL_016 vs. YAN_032
AUC: P = 0.0283
BEL_039 vs. YAN_014
AUC: P = 0.1283
BEL_005 vs. YAN_062
AUC: P = 0.1150
BEL_019 vs. YAN_001
AUC: P = 0.0163
BEL_010 vs. YAN_020
AUC: P = 0.2854
Serum glucose (mg/dl)
Time (min)
e
Colon – H&E
Ileum – H&E
Belgian
Yanomami
50 µm
f
Colon – AB/PAS
Colon mucus thickness (µm)
0.574
Belgian
Yanomami

***Suppl. Fig. 3|Degree of protection against HFD-induced metabolic pathology conferred by the gut microbiota varies among human donors.*** ***a***, *Weight follow-up of all (left), Belgian (middle) and Yanomami (right)-HMA mice over the course of high-fat diet (HFD) administration. Data are represented as mean ± SD.* ***b***, *Weight follow-up of separate donor sets over the course of high-fat diet (HFD) administration. Data are shown as mean ± SEM.* ***c***, *Blood glucose follow-up over the course of high-fat diet (HFD) administration. Data are represented as mean ± SD.* ***d,*** *Intraperitoneal glucose tolerance test (IPGTT). Mice were fasted for 16h and injected peritoneally with glucose (20%, 2g kg$^{-1}$ body weight for 1st experiment, 10%, 1g kg$^{-1}$ body weight for following experiments). Blood glucose levels were measured at the indicated time points. Data are shown as mean ± SD. Significance of area under the curve (AUC) was determined using a Welch's t test (P values indicated).* ***e***, *Representative colon (left) and ileum (right) histology images (H&E stain) of Belgian (above) vs. Yanomami (below) mice following 12 weeks of HFD.* ***f***, *Representative colon histology images (AB/PAS stain, left) of Belgian (above) vs. Yanomami (below) mice following 12 weeks of HFD and corresponding mucus thickness measurements based on AB/PAS-staining (right). Each datapoint represents a mouse, each symbol represents a different Belgian or Yanomami human microbiota. Mice with the same microbiota are treated as technical replicates. Significance was determined using a linear mixed-effects model fitted by REML (details in methods section, P value indicated).*

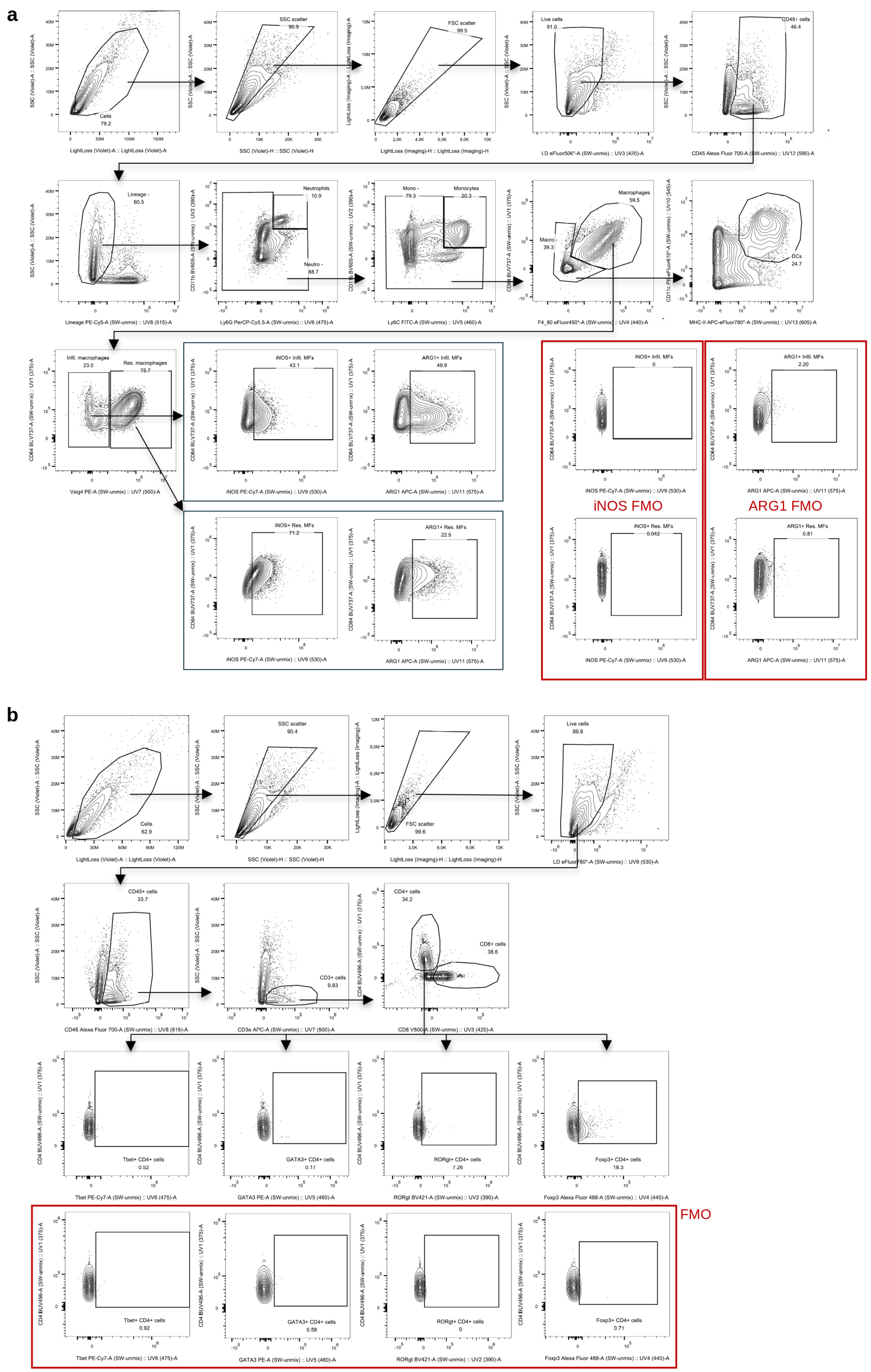


***Suppl. Fig. 4|Gating strategy used for immunophenotyping by flow cytometry. a,*** *Gating strategy used for the innate immune (myeloid) panel*

*to gate for neutrophils, monocytes, dendritic cells (DCs), macrophages (MFs), infiltrating (infil.) and resident (res.) macrophages, and iNOS/ARG1+ Infil./Res. MFs. Fluorescence Minus One (FMO) controls for iNOS and ARG1 are indicated by a red frame.* ***b,*** *Gating strategy used for T cell panel to gate for CD4/CD8+ T cells, and subsequently Th1/Th2/Th17/Tregs as Tbet+/GATA3+/RORγt+/Foxp3+ CD4+ T cells, respectively. FMO controls for Tbet/GATA3/RORγt/Foxp3 (in that order) are indicated by a red frame. Cell acquisition was done using the BD FACSDiscover A8 spectral cytometer, correcting for autofluorescence of the specific tissues.*

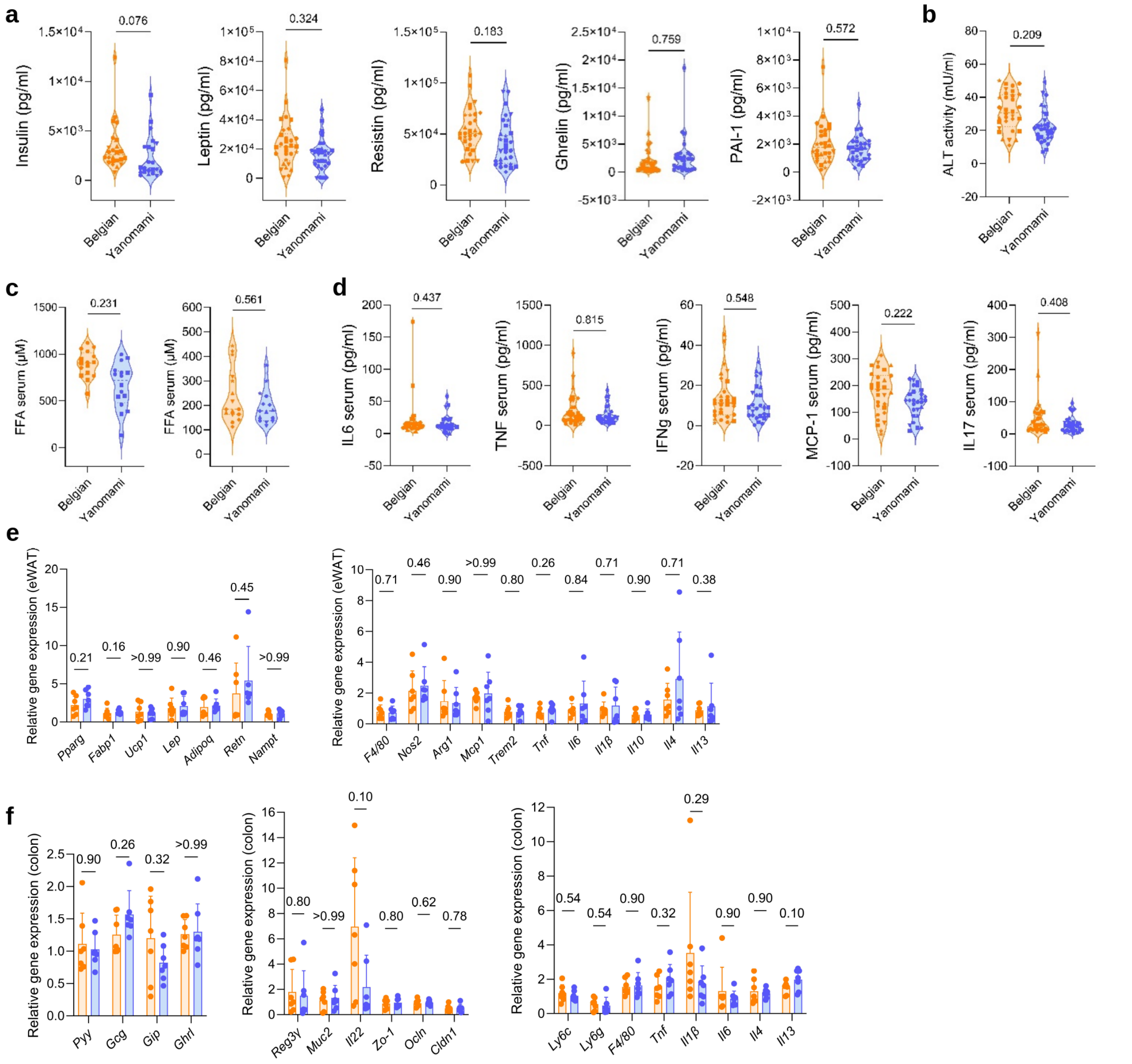


***Suppl. Fig. 5|Modest change in inflammatory and metabolic phenotype in Belgian vs. Yanomami-HMA mice upon HFD at the cohort level.*** ***a,*** *Diabetes markers measured in the fasting serum of mice following 12 weeks of HFD (16h fasting period). Each datapoint represents a mouse, each symbol represents a different Belgian or Yanomami human microbiota. Mice with the same microbiota are treated as technical replicates. For insulin and ghrelin, statistics was done on Log10 transformed data.* ***a-d****, Significance was determined using a linear mixed-effects model fitted by REML (details in methods section, P values indicated).* ***b****, Serum ALT activity measured in serum of mice following 12 weeks of HFD. Each datapoint represents a mouse, each symbol represents a different Belgian or Yanomami human microbiota. Mice with the same microbiota are treated as technical replicates.* ***c,*** *Free fatty acid (FFA) levels measured in serum of mice following 12 weeks of HFD. Results were split according to serum isolation circumstances, since this affected the read-out. Serum was isolated 120min post glucose injection for IPGTT for experiment comparing BEL_051 vs. YAN_009 and BEL_026 vs. YAN_013 (left), non-fasting serum was used for experiment with other donor sets (right). Each datapoint represents a mouse, each symbol represents a different Belgian or Yanomami human microbiota. Mice with the same microbiota are treated as technical replicates.* ***d,*** *Inflammatory cytokines measured in serum of mice following 12 weeks of HFD. Each datapoint represents a mouse, each symbol represents a different Belgian or Yanomami human microbiota. Mice with the same microbiota are treated as technical replicates. For IL6, statistics was done on Log10(x+1) transformed data; for IL17, statistics was done on Log10 transformed data.* ***e-f,*** *Gene expression analysis wherefor RNA was isolated from different tissue (eWAT and colon), and equal amounts of RNA per donor were pooled prior to RT-qPCR analysis.* ***e,*** *Gene expression analysis of markers related to metabolism and adipokines (left) or inflammation (right) in eWAT tissue of mice following 12 weeks of HFD. Gene expression was quantified relative to Gapdh and Hprt expression levels and scaled to a value of 1 for the first sample (Belgian 1). Significance was determined using a Mann-Whitney test (exact two-tailed P values indicated).* ***f,*** *Gene expression analysis of markers related to gut hormones (left), barrier function (middle) or inflammation (right) in colon tissue of mice following 12 weeks of HFD. Gene expression was quantified relative to Gapdh, Ubc and β-actin expression levels and scaled to a value of 1 for the first sample (Belgian 1). Significance was determined using a Mann-Whitney test (exact two-tailed P values indicated).*

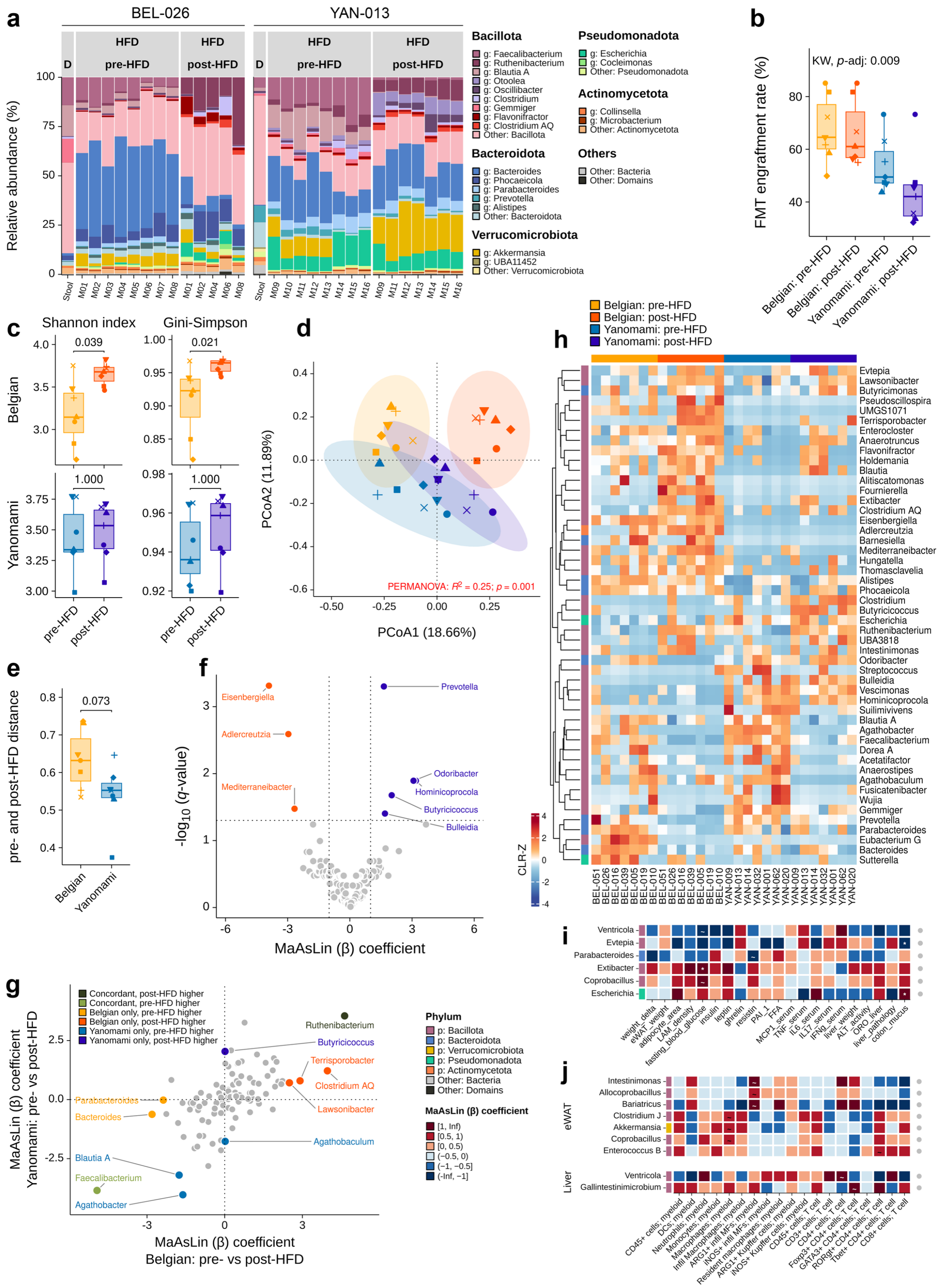

***Suppl. Fig. 6|Taxonomic composition, microbiota engraftment, and high-fat-diet responses across Belgian and Yanomami microbiota donors. a,*** *Genus-level relative abundance profiles from the main FMT experiment using Belgian donor BEL_026 (left) and Yanomami donor YAN_013 (right). The first bar represent donor stool (D), followed by individual pre- and post-HFD mouse samples. The ten most abundant Bacillota genera, five most abundant Bacteroidota genera, and three most abundant genera from each of Verrucomicrobiota, Pseudomonadota, and Actinomycetota are shown; remaining bacterial and non-bacterial taxa were grouped as "Other: Bacteria" and "Other: Domains", respectively. For* ***b–i****, one donor-level profile was used per human donor and time point (n = 7 per cohort and time point). For the BEL_051 versus YAN_009 and BEL_026 versus YAN_013 comparisons, profiles were averaged across individually sequenced mice; for all other donor comparisons, profiles generated from pooled samples were used directly.. Each symbol represents one human microbiota donor.* ***b,*** *Genus-level FMT engraftment, defined*

*as the proportion of donor-core genera accounting cumulatively for 99% of donor abundance that were detected in recipient mice profiles. Mean ± SD engraftment was 67.68 ± 12.70% and 66.17 ± 12.42% for Belgian pre- and post-HFD profiles, respectively, and 54.17 ± 10.59% and 44.28 ± 14.07% for Yanomami profiles (Kruskal–Wallis, P = 0.009).* ***c,*** *Donor-level Shannon and Gini–Simpson diversity before and after HFD. Diversity increased in Belgian-derived communities (Shannon: FDR-adjusted P = 0.039, estimated difference = −0.456, 95% CI −0.862 to −0.075; Gini–Simpson: adjusted P = 0.021, difference = −0.032, 95% CI −0.118 to −0.006), but not in Yanomami-derived communities (adjusted P = 1.000 for both indices; Wilcoxon rank-sum tests).* ***d,*** *PCoA of Bray–Curtis dissimilarities calculated from unaggregated taxonomic features. Community structure was associated with donor cohort and time point (PERMANOVA: $R^2$ = 0.255, F = 4.278, P = 0.001). Pairwise differences were detected between Belgian pre- versus post-HFD ($R^2$ = 0.299, P = 0.002), Belgian versus Yanomami pre-HFD ($R^2$ = 0.185, P = 0.002), Belgian pre-HFD versus Yanomami post-HFD ($R^2$ = 0.258, P = 0.001), Belgian post-HFD versus Yanomami pre-HFD ($R^2$ = 0.285, P = 0.001), Belgian versus Yanomami post-HFD ($R^2$ = 0.205, P = 0.002), and Yanomami pre- versus post-HFD profiles ($R^2$ = 0.178, P = 0.003). Within-group dispersion did not differ significantly (P = 0.088).* ***e,*** *Bray–Curtis dissimilarity between paired pre- and post-HFD donor-level profiles. Belgian-derived communities showed a slightly greater shift than Yanomami-derived communities (Wilcoxon rank-sum test, P = 0.073, estimated difference = 0.085, 95% CI −0.0037 to 0.193).* ***f,*** *Differentially abundant genera between Yanomami- and Belgian-derived post-HFD communities. Positive and negative MaAsLin2 coefficients (β) indicate associations with Yanomami- and Belgian-derived communities, respectively.* ***g,*** *Differential-abundance responses to HFD among genera detected in both groups. Genera associated with the same time point in both groups are highlighted in green; grey points indicate genera without significant associations. In* ***f–g****, significance was defined as |β| > 1 and BH-adjusted q < 0.05.* ***h,*** *Heatmap of the 50 genera with the greatest MaAsLin2 coefficients across contrasts. Rows were hierarchically clustered; annotations indicate phylum and combined donor cohort–time-point group.* ***i,*** *Associations between genera and host metabolic, inflammatory, and histopathological measurements after HFD.* ***j****, Associations between genera and flow-cytometry-derived immune-cell frequencies after HFD. Cell colours indicate MaAsLin2 coefficient direction and magnitude; asterisks indicate BH-adjusted q < 0.1. Values in parentheses indicate the numbers of Belgian (B) and Yanomami (Y) samples containing each genus, and the right annotation indicates phylum. Y-axes were ordered by hierarchical clustering using Euclidean distance and complete linkage to improve visualization. eWAT, epididymal white adipose tissue; FFA, free fatty acids; IL, interleukin; IFN-γ, interferon-γ; MCP-1, monocyte chemoattractant protein 1; PAI-1, plasminogen activator inhibitor 1; TNF, tumour necrosis factor.*

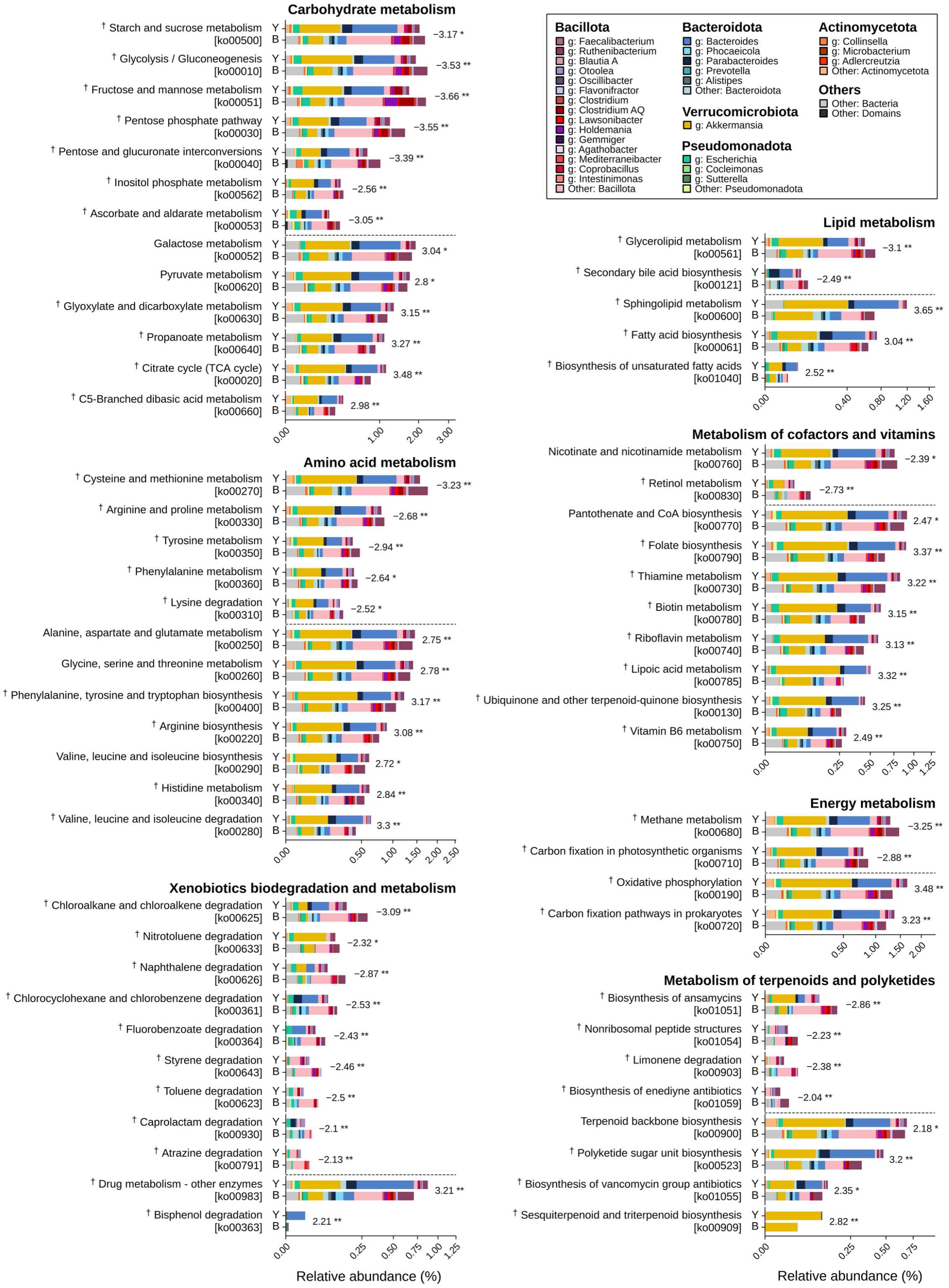


***Suppl. Fig. 7|Pathway differences between Belgian- and Yanomami-derived microbiota (BEL_026 vs. YAN_013) following high-fat diet.*** *Significantly associated KEGG metabolic pathways and genus-level contributions after HFD, grouped by KEGG metabolism subclass. Paired bars show mean pathway relative abundances in Belgian- and Yanomami-derived communities and are subdivided according to the genera contributing pathway-associated KOs. Signed values indicate LEfSe LDA scores, with positive and negative values denoting associations with Yanomami- and Belgian-derived communities, respectively. Significance was defined as |LDA| ≥ 2 and FDR-adjusted P < 0.05; ***adjusted P ≤ 0.001, **adjusted P ≤ 0.01, and *adjusted P ≤ 0.05. † indicates pathways independently supported by ANCOM-BC analysis at an adjusted P < 0.05. Square-root-scaled axes were used for visualization. The 15 most abundant Bacillota genera, five most abundant Bacteroidota genera, and three most abundant genera from each of Verrucomicrobiota, Pseudomonadota, and Actinomycetota are shown; remaining bacterial and non-bacterial contributions were grouped as “Other: Bacteria” and “Other: Domains”, respectively.*

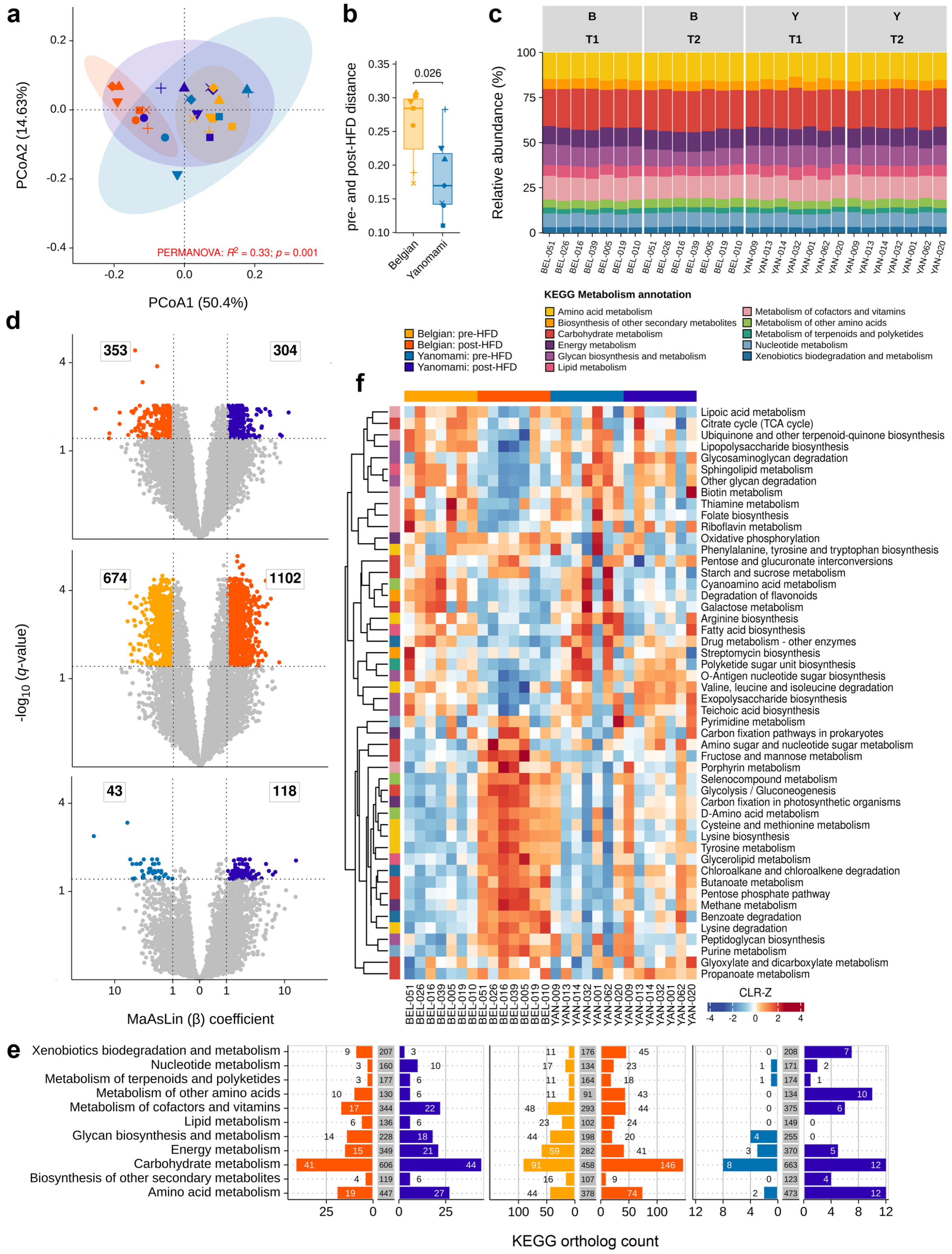


***Suppl. Fig. 8|Functional profiles and high-fat-diet responses across Belgian and Yanomami microbiota donors.*** *For all panels, one donor-level profile was used per human donor and time point (n = 7 per cohort and time point). For the BEL_051 versus YAN_009 and BEL_026 versus YAN_013 comparisons, profiles were averaged across individually sequenced mice; for all other donor comparisons, profiles generated from pooled samples were used directly. Each symbol represents one human microbiota donor. a, PCoA of Bray–Curtis dissimilarities calculated from KO profiles. Functional composition was associated with donor cohort and time point (PERMANOVA: $R^2$ = 0.326, F = 6.053, P = 0.001). Pairwise differences were detected between Belgian pre- and post-HFD profiles ($R^2$ = 0.617, P = 0.001), Belgian pre-HFD and Yanomami post-HFD profiles ($R^2$ = 0.218, P = 0.002), Belgian post-HFD and Yanomami pre-HFD profiles ($R^2$ = 0.446, P = 0.001), and Belgian and Yanomami post-HFD profiles ($R^2$ = 0.398, P = 0.003). No significant differences were detected between Belgian and Yanomami pre-HFD profiles ($R^2$ = 0.104, P = 0.189) or between Yanomami pre- and post-HFD profiles ($R^2$ = 0.126, P = 0.130). Within-group dispersion also differed (P = 0.017), indicating that both group centroids and dispersion may contribute to these differences. b, Bray–Curtis dissimilarity between paired pre- and post-HFD donor-level KO profiles. Belgian-derived communities showed a greater functional shift than Yanomami-derived communities (Wilcoxon rank-sum test, P = 0.026, estimated difference*

*= 0.0786, 95% CI 0.003–0.154). c, Relative abundance of KEGG metabolism subclasses before and after HFD, showing broadly conserved functional profiles across groups. d, Differentially abundant KOs for Belgian versus Yanomami post-HFD profiles (top), Belgian pre- versus post-HFD profiles (centre), and Yanomami pre- versus post-HFD profiles (bottom). Significance was defined as |MaAsLin2 coefficient (β)| > 1 and BH-adjusted q < 0.05. In the post-HFD population comparison, 353 KOs were associated with Belgian-derived communities and 304 with Yanomami-derived communities. Within Belgian-derived communities, 674 KOs were associated with pre-HFD and 1,102 with post-HFD profiles, whereas within Yanomami-derived communities, 43 KOs were associated with pre-HFD and 118 with post-HFD profiles. Square-root-scaled axes were used for visualization. e, Distribution of differentially abundant KO assignments across KEGG metabolism subclasses for the contrasts shown in d. Labels indicate the number associated with each population or time point; central grey bars represent non-differentially abundant KOs. f, Heatmap of the 50 metabolic pathways with the greatest absolute MaAsLin2 coefficients across contrasts. Rows were hierarchically clustered; annotations indicate KEGG metabolism subclass and combined donor cohort–time-point group.*

### Supplementary tables

***Suppl. Table 1|Antibodies used for flow cytometry analysis.***

| Marker | Fluorophore | Clone | Supplier | Catalog # |
|---|---|---|---|---|
| LiveDead | eFluor506 | | eBioscience | 65-0866-14 |
| Livedead | efluor780 | | eBioscience | 65-0865-14 |
| CD64 | Biotin | X54-5/7.1 | BioLegend | 139318 |
| F4/80 | eFluor450 | BM8 | Invitrogen | 48-4801-82 |
| Ly-6G | PerCP-Cy5.5 | 1A8 | BD | 560602 |
| SAV | BUV737 | | BD | 612775 |
| CD11b | BV605 | M1/70 | BD | 563015 |
| Ly-6C | FITC | AL-21 | BD | 553104 |
| CD45 | AF700 | 30-F11 | Invitrogen | 56-0451-82 |
| MHC-II | APC-eFluor780 | M5/114.15.2 | Invitrogen | 47-5321-80 |
| CD11c | PE-eFluor610 | N418 | Invitrogen | 61-0114-82 |
| CD3e | PE-Cy5 | 145-2C11 | Invitrogen | 15-9031-82 |
| CD19 | PE-Cy5 | 1D3 | Invitrogen | 15-0193-82 |
| NK1.1 | PE-Cy5 | PK136 | BioLegend | 108716 |
| Vsig4 | PE | NLA14 | Invitrogen | 12-5752-82 |
| iNOS | PE-Cy7 | CXNFT | Invitrogen | 25-5920-80 |
| ARG1 | APC | A1exF5 | Invitrogen | 17-3697-80 |
| CD3e | APC | 145-2C11 | Invitrogen | 17-0031-83 |
| CD4 | BUV496 | GK1.5 | BD | 612952 |
| CD8 | V500 | 53-6.7 | BD | 560776 |
| Foxp3 | AF-488 | FJK-16s | Invitrogen | 53-5773-82 |
| GATA3 | PE | TWAJ | Invitrogen | 12-9966-42 |
| RORgt | BV421 | Q31-378 | BD | 562894 |
| Tbet | PE-Cy7 | eBio4B10 | Invitrogen | 25-5825-82 |

***Suppl. Table 2|Primers used for gene expression analysis (RT-qPCR).***

| Gene | Forward sequence (5' – 3') | Reverse sequence (5' – 3') |
|---|---|---|
| *Hprt* | AGTGTTGGATACAGGCCAGAC | CGTGATTCAAATCCCTGAAGT |
| *Ubc* | AGGTCAAACAGGAAGACAGACGTA | TCACACCCAAGAACAAGCACA |
| *Gapdh* | TGAAGCAGGCATCTGAGGG | CGAAGGTGGAAGAGTGGGAG |
| *β-actin* | GCTTCTAGGCGGACTGTTACTGA | GCCATGCCAATGTTGTCTCTTAT |
| *Ywhaz* | GGAAAGTTCTTGATCCCCAATGC | TGTGACTGGTCCACAATTCCTT |
| *Rplp0* | AGATTCGGGATATGCTGTTGGC | TCGCCTCCTAGACCAGTGTTC |
| *Trem2* | CTACCAGTGTCAGAGTCTCCGA | CCTCGAAACTCGATGACTCCTC |
| *Tnf* | ACCCTGGTATGAGCCCATATAC | ACACCCATTCCCTTCACAGAG |
| *F4/80* | GATGCCCAAGGATCTGAAAA | CTCTGTGGTCCCACCTTCAT |
| *Mcp1* | AGCTCTCTCTTCCTCCACCAC | TGCTGGTGATCCTCTTGTAGC |
| *Il6* | CCGGAGAGGAGACTTCACAG | TTCTGCAAGTGCATCATCGT |
| *Il1β* | GAAATGCCACCTTTTGACAGTG | TGGATGCTCTCATCAGGACAG |
| *Il10* | GCCTTATCGGAAATGATCCA | TTTTCACAGGGGAGAAATCG |
| *Il4* | TGGACTCATTCATGGTGCAG | AACATGGGAAAACTCCATGC |
| *Il13* | TGTGTAGCTGAGCAGTTTTGT | TTCTGTGTAGCCCTGGATTCC |
| *Ly6c* | GCAGTGCTACGAGTGCTATGG | ACTGACGGGTCTTTAGTTTCCTT |
| *Ly6g* | TGGACTCTCACAGAAGCAAAG | GCAGAGGTCTTCCTTCCAACA |
| *Nos2* | CAGCTGGGCTGTACAAACCTT | CATTGGAAGTGAAGCGTTTCG |
| *Arg1* | AAAGCTGGTCTGTGGAAAA | ACAGACCGTGGGTTCTTCAC |
| *Pparγ* | AGACCACTCGCATTCCTTTG | TCGCACTTTGGTATTCTTGG |
| *Fabp4* | TCACCATCCGGTCAGAGAGTA | TCCTGTCGTCTGCGGTGATT |
| *Leptin* | CAGGGAGGAAAATGTGCTGGAG | CCGACTGCGTGTCTGAAATGTC |
| *Adipoq* | GAAGCCGCTTATGTGTATCGC | GAATGGGTACATTGGGAACAGT |
| *Retn* | GTACCCACGGGATGAAGAACC | GCAGAGCCACAGGAGCAG |
| *Nampt* | GGCACCACTAATCATCAGACCTG | AAGGTGGCAGCAACTTGTAGCC |
| *Ucp1* | ACTGCCACACCTCCAGTCATT | CTTTGCCTCACTCAGGATTGG |
| *Pyy* | GCAGCGGTATGGAAAAAGAGA | TCCAAACCTTCTGGCCTGAA |
| *Gcg* | CCTTCAAGACACAGAGGAGAACC | CTGTAGTCGCTGGTGAATGTGC |
| *Gip* | CAAGACTTCGTGAACTGGCTGC | GTCCTCCTTTCCCTGAGATTGC |
| *Ghrl* | GAAGCCACCAGCTAAACTGCAG | CTGACAGCTTGATGCCAACATCG |
| *Reg3γ* | CCTCAGGACATCTTGTGTCTGTGCTC | TCCACCTCTGTTGGGTTCATAGCC |
| *Muc2* | TCCCAGACAGGTTAATGAGAC | CAGCAGTTATCAGCATCAGG |
| *Il22* | ATCAGTGCTACCTGATGAAG | CATTCTTCTGGATGTTCTGG |
| *Zo-1* | CCACCTCTGTCCAGCTCTTC | CACCGGAGTGATGGTTTTCT |
| *Ocln* | TTGAAAGTCCACCTCCTTACAGA | CCGGATAAAAAGAGTACGCTGG |
| *Cldn1* | TGCCCCAGTGGAAGATTTACT | CTTTGCGAAACGCAGGACAT |

***Suppl. Table 3|Sequencing, assembly and annotation quality control for donor-specific mouse faecal metagenome co-assemblies.*** *Mouse faecal metagenomic reads were preprocessed and co-assembled separately for each human microbiota donor. Raw reads and reads after quality control (QC) are shown in millions as mean ± SD across sequenced samples contributing to each co-assembly, where applicable. Reads after QC were retained following quality filtering and removal of host and technical sequences. High-quality read mapping rate indicates the mean ± SD percentage of filtered reads mapping back to the corresponding co-assembly with mapping quality ≥20. Assembly-level metrics, reported once per donor-specific co-assembly, comprise contig number, total assembly length, contig N50, predicted protein-coding sequences (CDS), the percentage of predicted CDS assigned at least one Kyoto Encyclopedia of Genes and Genomes (KEGG) Orthology (KO) identifier, and the percentages of assembled contigs assigned taxonomically at least to phylum or genus level. CDS were predicted using Prokka, KO annotations were assigned using eggNOG-mapper against eggNOG, and contig taxonomy was assigned using MMseqs2 against the Genome Taxonomy Database (GTDB). The final column reports mean ± SD across all co-assemblies.*

| Attribute | BEL_051 | YAN_009 | BEL_026 | YAN_013 | BEL_016 | YAN_032 | BEL_039 | YAN_014 | BEL_005 | YAN_062 | BEL_019 | YAN_001 | BEL_010 | YAN_020 | Overall average |
|---|---|---|---|---|---|---|---|---|---|---|---|---|---|---|---|
| Raw reads (million) | 31.81 (±10.80) | 31.17 (±9.61) | 23.95 (±3.62) | 36.13 (±30.10) | 20.11 (±0.95) | 20.88 (±1.00) | 17.56 (±1.38) | 18.97 (±2.77) | 20.13 (±0.08) | 20.35 (±1.86) | 19.17 (±0.46) | 21.87 (±0.89) | 19.28 (±1.86) | 20.09 (±0.30) | 22.96 (±4.69) |
| Reads after QC (million) | 30.92 (±10.37) | 25.93 (±5.77) | 21.37 (±5.90) | 33.28 (±26.19) | 19.69 (±1.32) | 20.11 (±1.97) | 16.93 (±0.56) | 18.10 (±1.66) | 19.75 (±0.44) | 19.61 (±2.80) | 18.99 (±0.47) | 21.14 (±0.01) | 17.32 (±4.32) | 19.35 (±0.55) | 21.61 (±4.45) |
| Contigs (×10³) | 185.14 | 173.31 | 173.94 | 66.45 | 74.82 | 51.67 | 104.30 | 65.47 | 116.47 | 54.42 | 95.40 | 74.75 | 90.40 | 97.98 | 101.75 (±47.67) |
| Assembly length (Mbp) | 730.80 | 529.02 | 619.90 | 302.04 | 256.13 | 193.88 | 265.79 | 180.50 | 317.46 | 180.05 | 241.97 | 209.27 | 267.23 | 229.35 | 323.10 (±174.07) |
| Contig N50 (kbp) | 27.50 | 14.11 | 26.74 | 35.68 | 18.51 | 26.20 | 7.62 | 20.71 | 9.06 | 20.38 | 9.43 | 12.32 | 17.29 | 6.47 | 18.00 (±8.78) |
| HQ read mapping rate (%) | 89 (±4.3) | 89.6 (±1.5) | 94.1 (±1.5) | 92 (±3) | 87.9 (±1.3) | 89.4 (±3.3) | 94 (±0) | 93.8 (±0.1) | 92.2 (±0.2) | 90.3 (±0.2) | 89.7 (±1.8) | 93.2 (±0) | 87.9 (±1.1) | 89.7 (±0.4) | 90.9 (±1.5) |
| Predicted CDS (×10³) | 691.29 | 508.20 | 585.33 | 285.97 | 239.32 | 178.24 | 250.27 | 172.98 | 297.40 | 169.06 | 226.98 | 196.78 | 250.28 | 218.79 | 305.06 (±165.72) |
| KO annotation rate (%) | 64.23 | 62.52 | 63.41 | 63.37 | 62.35 | 60.97 | 59.88 | 59.92 | 62.87 | 62.59 | 61.84 | 60.93 | 63.13 | 61.30 | 62.09 (±1.33) |
| Phylum-level annotation rate (%) | 94.76 | 94.23 | 93.15 | 94.81 | 94.21 | 93.84 | 95.64 | 95.32 | 95.38 | 95.24 | 95.08 | 93.41 | 94.93 | 94.78 | 94.63 (±0.76) |
| Genus-level annotation rate (%) | 70.52 | 70.95 | 70.34 | 73.54 | 69.29 | 70.60 | 77.75 | 72.66 | 76.70 | 70.04 | 76.76 | 73.46 | 75.82 | 71.74 | 72.87 (±2.86) |
